# Agonist binding affinity determines palmitoylation of the glucagon-like peptide-1 receptor and its functional interaction with plasma membrane nanodomains in pancreatic beta cells

**DOI:** 10.1101/492496

**Authors:** Teresa Buenaventura, William E. Laughlin, Stavroula Bitsi, Thomas Burgoyne, Zekun Lyu, Affiong I. Oqua, Hannah Norman, Emma R. McGlone, Andrey S. Klymchenko, Ivan R Corrêa, Abigail Walker, Asuka Inoue, Aylin Hanyaloglu, Guy A. Rutter, Stephen R. Bloom, Ben Jones, Alejandra Tomas

## Abstract

The glucagon-like peptide-1 receptor (GLP-1R), a key pharmacological target in type 2 diabetes and obesity, is known to undergo palmitoylation by covalent ligation of an acyl chain to cysteine 438 in its carboxyl-terminal tail. Work with other GPCRs indicates that palmitoylation can be dynamically regulated to allow receptors to partition into plasma membrane nanodomains that act as signaling hotspots. Here, we demonstrate that the palmitoylated state of the GLP-1R is increased by agonist binding, leading to its segregation and clustering into plasma membrane signaling nanodomains before undergoing internalization in a clathrin-dependent manner. Both GLP-1R signaling and trafficking are modulated by strategies targeting nanodomain segregation and cluster formation, including depletion of cholesterol or expression of a palmitoylation-defective GLP-1R mutant. Differences in receptor binding affinity exhibited by biased GLP-1R agonists, and modulation of binding kinetics with the positive allosteric modulator BETP, influence GLP-1R palmitoylation, clustering, nanodomain signaling, and internalization. Downstream effects on insulin secretion from pancreatic beta cells indicate that these processes are relevant to GLP-1R physiological actions and might be therapeutically targetable.

## Introduction

G protein-coupled receptors (GPCRs), the largest membrane receptor family in eukaryotes (1), are integral membrane proteins and, as such, both their physical organization and their signaling properties are modulated by the lipid composition of the surrounding membrane (2, 3). The localization of GPCRs to dynamic membrane nanodomains has been widely reported (4-6). These nanodomains, or membrane rafts, which cannot be directly observed in living cells with current methods (7), are often described as highly organized detergent-resistant, liquid-ordered, glycosphingolipid-and cholesterol-rich platforms where receptor-signaling complexes become compartmentalized, facilitating efficient coupling with G proteins (5, 8, 9). Additionally, most GPCRs are modified post-translationally with one or more palmitic acid chains linked covalently, but reversibly, via a thioester bond to cysteines within the intracellular domain of the receptor, in a process known as palmitoylation (2, 10). The insertion of acyl chain(s) is regulated by families of acyltransferases (DHHCs) and palmitoyl protein thioesterases (11, 12), and can be either constitutive or modulated by agonist binding (13, 14). GPCR palmitoylation at or near the end of its carboxyl-terminal (C-terminal) tail creates a new membrane anchor and a further intracellular loop (15) that modifies the GPCR structure and its interactions with specific intracellular partners, favoring receptor partitioning into plasma membrane nanodomains (2, 16, 17).

We have previously described how signaling responses of the glucagon-like peptide-1 receptor (GLP-1R), a class B GPCR and prime type 2 diabetes target, are modulated by intracellular membrane trafficking processes (18, 19). In the present study, we establish that palmitoylation of the GLP-1R, known to occur at cysteine 438 near the C-terminal end of its cytoplasmic tail (20), is significantly increased after binding to the pharmacological agonist exendin-4, and leads to GLP-1R clustering into cholesterol-rich plasma membrane nanodomains that enable receptor signaling and endocytosis. Surprisingly, although β-arrestin-2 recruitment to the receptor is enhanced within detergent-resistant membrane fractions, indicating that active GLP-1Rs coalesce in these liquid-ordered nanodomains, β-arrestins were dispensable for GLP-1R internalization. Instead, the presence of cholesterol is the main requirement for effective GLP-1R signaling and internalization, indicating that plasma membrane compartmentalization underlies GLP-1R action.

Given that the palmitoylation of GPCRs can regulate biased agonist signaling due to the control of their redistribution into membrane nanodomains (21, 22), we have also determined the effect of biased GLP-1R agonists derived from exendin-4 previously characterized by us for their differing efficacy as glucose lowering agents *in vivo* (19), on agonist-induced GLP-1R palmitoylation, clustering and nanodomain signaling. We find that GLP-1R palmitoylation is potentiated by a biased agonist that exhibits high receptor binding affinity, which correlates with increased β-arrestin recruitment and GLP-1R internalization, as well as higher levels of GLP-1R clustering and signaling from liquid-ordered nanodomains compared to the parental peptide. Conversely, an agonist biased in the opposite direction, with reduced receptor binding affinity, β-arrestin recruitment and GLP-1R internalization, triggers reduced levels of GLP-1R palmitoylation, clustering and signaling from liquid-ordered nanodomains. The reduced GLP-1R nanodomain partitioning, signaling and internalization of the low-affinity biased agonist is, however, restored by allosterically increasing its residence time at the receptor using the GLP-1R-specific positive allosteric modulator (PAM) 2-(Ethylsulfinyl)-4-[3-(phenylmethoxy)phenyl]-6-(trifluoromethyl)-pyrimidine (BETP) (23, 24).

This is, to our knowledge, the first report linking biased agonist binding affinities to the control of receptor palmitoylation and nanodomain partitioning to modified responses of a class B GPCR, opening the door for future studies based on the direct manipulation of these processes to generate new tools for the control GLP-1R action with the potential to translate into more effective diabetes therapies.

## Results

Prompted by a previous report demonstrating palmitoylation at cysteine 438 of the cytoplasmic tail of the GLP-1R (20), we analyzed whether this post-translational modification is constitutive or induced by agonist binding. Using Chinese Hamster Ovary (CHO)-K1 cells stably expressing human GLP-1R SNAP-tagged at the extracellular N-terminus (19, 25), we detected a very low level of basal GLP-1R palmitoylation that was significantly increased following stimulation with the orthosteric peptide agonist exendin-4 (Figure 1A) in glycosylated GLP-1Rs (Supplementary Figure 1A), known to represent the functional pool of the receptor for agonist-binding (26) at the cell surface (Supplementary Figure 1B). To confirm the relevance and specificity of these results to pancreatic beta cells, we repeated the palmitoylation assay in MIN6B1 mouse beta cells (27) stably expressing SNAP-GLP-1R (18) in the presence or absence of the palmitoylation inhibitor 2-bromopalmitate (Supplementary Figure 1C).

**Figure 1.**
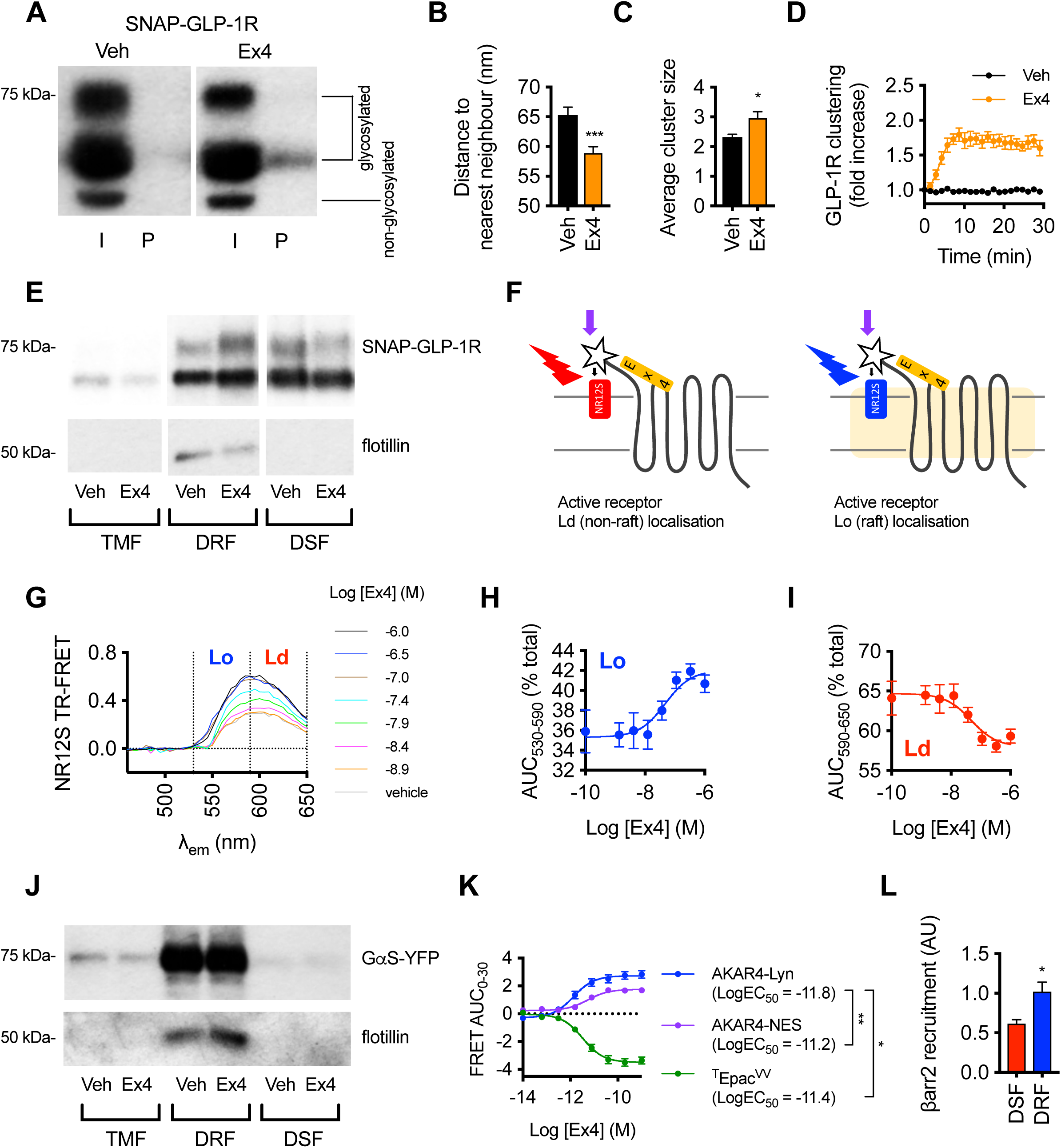
Agonist-induced SNAP-GLP-1R palmitoylation, clustering and recruitment to membrane nanodomains. (**A**) Total input (I) and palmitoylated (P) SNAP-GLP-1R fractions from CHO-K1 cells stably-expressing SNAP-GLP-1R and treated with vehicle (Veh) or 100 nM exendin-4 (Ex4) for 10 min. (**B**) Distance to the nearest neighbor quantified by EM analysis of 2-D plasma membrane sheets isolated from MIN6B1 beta cells stably-expressing SNAP-GLP-1R following gold labeling of SNAP-tagged receptors and treatment with vehicle or 100 nM exendin-4 for 2 min, data is from a minimum of *n*=1,000 gold particles per condition, unpaired *t*-test. (**C**) Average cluster sizes calculated from TIRF-PALM measurements of HEK293 cells stably expressing FLAG-GLP-1R and labelled with anti-FLAG CAGE 500 antibodies prior to stimulation with vehicle or 100 nM exendin-4 for 2 min, data from *n*=11 regions per condition, paired *t*-test. (**D**) Agonist-induced SNAP-GLP-1R clustering in monoclonal CHO-SNAP-GLP-1R cells dual-labeled with Lumi4-Tb and SNAP-Surface 647, treated with 100 nM exendin-4 or vehicle, TR-FRET displayed as fold increase relative to individual baseline, *n*=6. (**E**) SNAP-GLP-1R distribution within total membrane (TMF), detergent-resistant (DRF) and detergent-soluble (DSF) fractions isolated from MIN6B1 cells stably-expressing SNAP-GLP-1R treated with vehicle or 100 nM exendin-4 for 2 min, with flotillin as a marker of membrane raft enrichment, quantification shown in Supplementary Figure 1E. (**F**) Cartoon explaining NR12S assay to monitor SNAP-GLP-1R translocation to membrane nanodomains. Ld, liquid-disordered; Lo, liquid-ordered. (**G**) NR12S-associated TR-FRET spectra in Lumi4-labeled HEK293 cells stably expressing SNAP-GLP-1R treated with the indicated concentrations of exendin-4 or vehicle, Lumi4-Tb-only spectrum has been subtracted, and spectrum divided into Lo (530 - 590 nm) and Ld (590 - 650 nm) regions, *n*=6, error bars not shown for clarity. (**H**) Preferential increase in Lo-associated SNAP-GLP-1R-NR12S TR-FRET induced by exendin-4, determined from (G) as percentage of total AUC from 530 - 590 nm portion of the spectrum, 3-parameter fit shown. (**I**) Decrease in Ld-associated SNAP-GLP-1R-NR12S TR-FRET induced by exendin-4, determined from (G) as percentage of total AUC from 590 - 650 nm portion of the spectrum, 3-parameter fit shown. (**J**) GαS-YFP distribution within total membrane (TMF), detergent-resistant (DRF) and detergent-soluble (DSF) fractions isolated from MIN6B1 cells stably expressing SNAP-GLP-1R and transiently transfected with GαS-YFP prior to treatment with vehicle or 100 nM exendin-4 for 2 min, with flotillin as a marker of membrane raft enrichment. (**K**) Dose-responses for exendin-4-induced FRET changes indicating PKA activation within membrane rafts (AKAR4-Lyn, *n*=11), or cytosol (AKAR4-NES, *n*=5), as well as total cellular cAMP (^T^Epac^VV^, *n*=15), in CHO-SNAP-GLP-1R cells, determined by calculation of AUC of baseline-normalized FRET ratio from sequential measurements over 30 min, 3-parameter fit, FRET traces shown in Supplementary Fig. 1H-J. Potency values (Log EC_50_) compared by one-way ANOVA with Tukey’s test. (**L**) Level of β-arrestin-2 recruitment to the GLP-1R in detergent-soluble (DSF) *vs.* detergent-resistant (DRF) fractions of CHO-PathHunter (DiscoverX) cells expressing ProLink-tagged human GLP-1R and β-arrestin-2-EA. β-arrestin-2 recruitment calculated for exendin-4-treated over vehicle conditions, and normalized to GLP-1R levels for each membrane fraction and to level of β-arrestin-2 recruitment of the total membrane fraction (TMF) for each experimental repeat, *n*=4 experiments, paired *t*-test. Unless stipulated, data are depicted as mean ± SEM; *p<0.05, **p<0.01, ***p<0.001 by statistical test indicated in the text.

As palmitoylation is known to regulate clustering of cell surface receptors, we next determined the degree of GLP-1R clustering at the plasma membrane before and after stimulation with exendin-4. We performed electron microscopy (EM) analysis of intact 2-D sheets of apical membranes ripped-off from adherent MIN6B1 cells stably expressing SNAP-GLP-1R and labeled live with the membrane-impermeable BG-SS-PEG4-biotin SNAP-tag probe followed by gold-conjugated streptavidin. Quantification of gold particle distances to the nearest neighbor showed increased clustering of SNAP-GLP-1Rs following exendin-4 stimulation (Figure 1B, Supplementary Figure 1E). These results were supported by total internal reflection fluorescence (TIRF)-photoactivatable localization microscopy (PALM) analysis of HEK293 cells stably expressing FLAG-GLP-1R labeled live with a photo-activatable caged dye, a technique that allows localization of receptors at the cell membrane with an accuracy of ∼10nm (28). We detected an increase in the average number of receptors per cluster after 1-minute stimulation with exendin-4 (Figure 1C). Note that receptor oligomers were present in basal conditions, in keeping with a previous report (29).

To gain more kinetic information on agonist-induced GLP-1R clustering, we next used a dual surface labeling approach that allows detection of receptor-receptor interactions by time-resolved (TR)-Förster resonance energy transfer (FRET). Due to the long-lived fluorescence of the lanthanide FRET donor (Lumi4-Tb), energy transfer results from both stable and transient protein-protein interactions, which increases when receptors are in closer proximity due to lateral diffusion within the membrane (30, 31). In CHO-K1 cells stably expressing SNAP-GLP-1R, 100 nM exendin-4 induced a rapid increase in TR-FRET with a t_1/2_ of 3.7±0.3 minutes (Figure 1D).

Increased clustering might reflect increased intermolecular proximity following receptor segregation into detergent-resistant dynamic membrane nanodomains (8). To investigate this possibility, we measured SNAP-GLP-1R levels in purified detergent-resistant and detergent-soluble fractions from total membrane preparations of MIN6B1 cells stably expressing SNAP-GLP-1R. Stimulation with exendin-4 resulted in an increased recruitment of the receptor to the flotillin-positive detergent-resistant fraction compared to vehicle control conditions, suggesting increased partitioning of SNAP-GLP-1R into membrane nanodomains (Figure 1E, quantified in Supplementary Figure 1F). Nanodomain partitioning in response to exendin-4 was confirmed in HEK293 cells expressing SNAP-GLP-1R (Supplementary Figure 1G). To corroborate this biochemical finding, we also developed a new, simple, medium-throughput method to monitor SNAP-GLP-1R localization in cholesterol-rich nanodomains using TR-FRET and the solvatochromic membrane probe NR12S (32). This Nile Red-derived probe partitions similarly into the outer membrane leaflet of both liquid-ordered and liquid-disordered membrane phases, but its spectral properties depend on the lipidic environment in which it is situated (Figure 1F), i.e. its fluorescence emission is blue-shifted in liquid-ordered compared to liquid-disordered membrane environments. In HEK293 cells expressing SNAP-GLP-1R labeled with Lumi4-Tb, TR-FRET was detectable upon addition of NR12S at several concentrations, which was further increased on addition of exendin-4 (Supplementary Figure 1H,I). The latter can be explained by the relative movement between the Lumi4-Tb-labeled extracellular domain of the SNAP-GLP-1R and the plasma membrane upon agonist binding, i.e. the assay senses ligand binding-triggered conformational rearrangements of receptors, as previously demonstrated with the epidermal growth factor (EGF) receptor (33). However, the exendin-4-induced TR-FRET increase was preferentially seen in the part of the NR12S spectrum associated with liquid-ordered localizations, in a dose-dependent manner, with a corresponding proportional decrease (relative to total) in liquid-disordered-associated signal (Figure 1G-I), matching our earlier observation that exendin-4 stimulation causes SNAP-GLP-1R translocation into detergent-resistant membrane nanodomains. Similar results were obtained when data were reanalyzed with single wavelengths assigned to liquid-ordered (570 nm) and liquid-disordered (610 nm) phases, reducing the requirement for full spectral measurements (Supplementary Figure 1J). The assay was also adapted to CHO-K1 cells stably expressing SNAP-GLP-1R (Supplementary Figure 1K,L).

Membrane nanodomains have often been described as hotspots for signaling as a result of co-segregation of membrane receptors with their corresponding signaling effectors (5). In order to test whether this is the case for SNAP-GLP-1R-enriched plasma membrane nanodomains in pancreatic beta cells, we purified detergent-resistant and detergent-soluble fractions from MIN6B1 cells transfected with a vector expressing the Gα_S_ subunit, the primary signaling effector coupled to the GLP-1R (34), fused to YFP. We found that Gα_S_-YFP clearly partitioned into detergent-resistant fractions, both before and after GLP-1R stimulation with exendin-4 (Figure 1J). In keeping with the distribution pattern of Gα_S_ at the plasma membrane, we also found that exendin-4 preferentially stimulated the cAMP-dependent GLP-1R downstream effector protein kinase A (PKA) within liquid-ordered nanodomains, as determined by comparing the activation potencies of the FRET-based conformational biosensors AKAR4-Lyn [PKA sensor targeted to cholesterol-rich nanodomains (35)], AKAR4-NES (cytosolic PKA sensor) and ^T^Epac^VV^ [total cAMP sensor (36)] (Figure 1K; Supplementary Figure 1M-O). Moreover, following exendin-4 stimulation, recruitment of the active GPCR regulatory factor β-arrestin-2 (37) to the GLP-1R was significantly increased in detergent-resistant *versus* detergent-soluble fractions of CHO-PathHunter (DiscoverX) cells expressing ProLink-tagged human GLP-1R and β-arrestin-2-EA (Figure 1L).

In comparison, and highlighting that agonist-dependent modulation of palmitoylation and recruitment to membrane nanodomains are receptor-specific responses, the closely related class B GPCR glucose-dependent insulinotropic polypeptide (GIP) receptor (GIPR) displayed a high degree of constitutive palmitoylation but no GIP-induced increases in either palmitoylation or recruitment to detergent-resistant fractions (Supplementary Figure 2A,B). Correspondingly, constitutive clustering of SNAP-GIPR in transiently transfected HEK293 cells was higher than that observed for SNAP-GLP-1R, despite similar receptor surface expression levels, with no detectable increase upon agonist addition (Supplementary Figure 2C).

**Figure 2.**
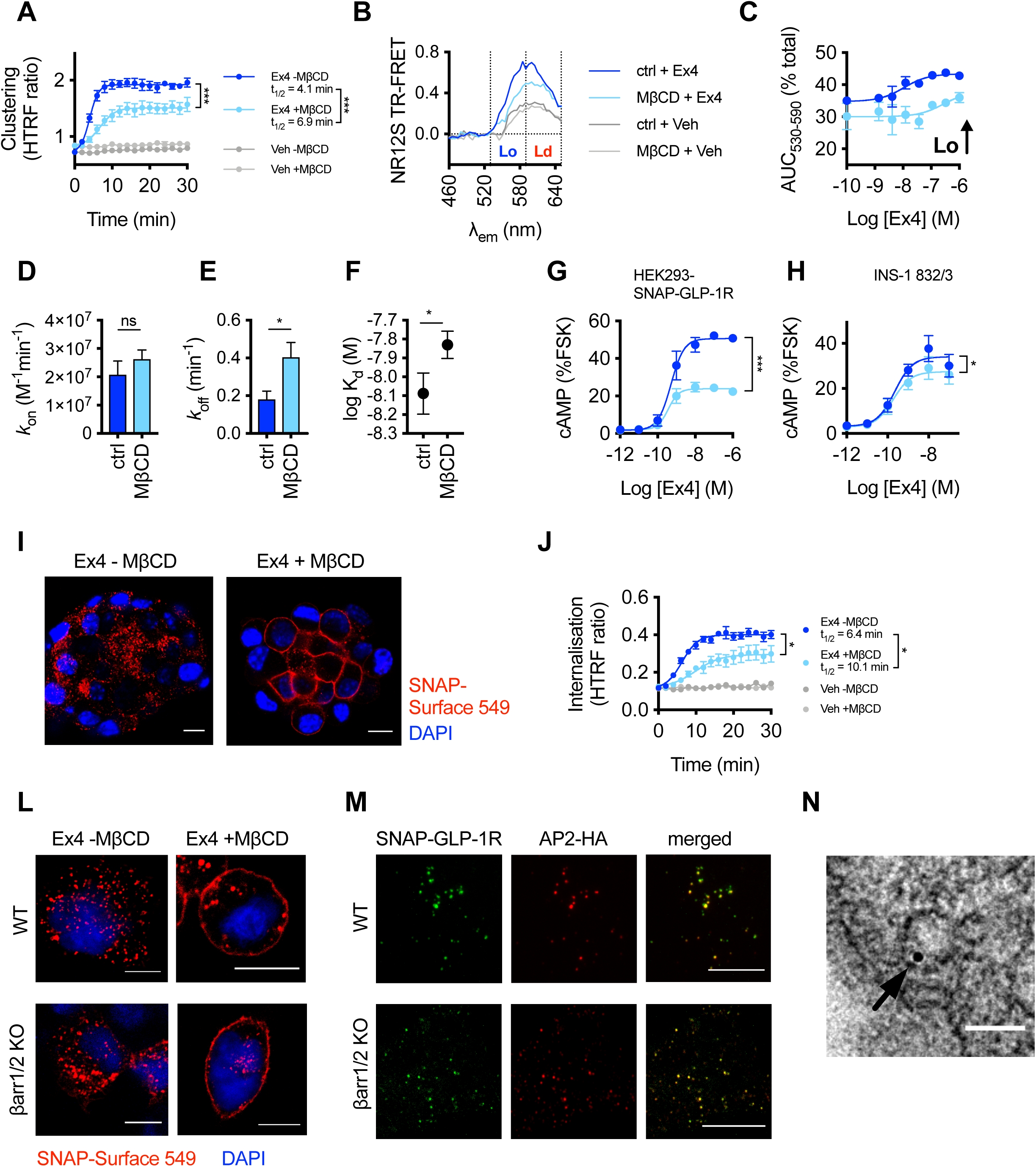
Effects of cholesterol depletion by MβCD treatment on clustering, membrane nanodomain recruitment, signaling and internalization properties of SNAP-GLP-1Rs. (**A**) SNAP-GLP-1R clustering measured by HTRF in HEK293 cells stably-expressing SNAP-GLP-1R with and without prior cholesterol depletion with MβCD (10 mM, 45 min), treated with exendin-4 (Ex4, 100 nM) or vehicle (Veh), expressed as HTRF ratio, *n*=4, 4-parameter logistic fit shown and used to quantify E_max_ and t_1/2_ (compared by paired *t*-tests). (**B**) NR12S TR-FRET spectra with and without cholesterol depletion in HEK-SNAP-GLP-1R cells by MβCD (10 mM), with either vehicle or exendin-4 (1 µM) treatment, shown after subtraction of the Lumi4-Tb response, *n*=3 parallel experiments, error bars omitted for clarity. (**C**) Dose-response analysis of MβCD effect on exendin-4-induced Lo-associated SNAP-GLP-1R-NR12S TR-FRET, determined from the same experiment as in (B), calculated as percentage of total AUC from 530 - 590 nm portion of spectrum, 3-parameter fit shown. (**D**) Association rate constant (*k*_on_) determined from TR-FRET kinetic binding response of exendin-4-FITC measured in Lumi4-Tb-labeled HEK-SNAP-GLP-1R cells, with or without treatment with MβCD (10 mM, 45 min), expressed ratiometrically as 520 / 620 nm signal after blank subtraction, *n*=4, non-significant (ns), paired *t*-test. TR-FRET traces shown in Supplementary Figure 2E. (**E**) Dissociation rate constant (*k*_off_) determined from same dataset as (D), paired *t*-test. (**F**) Equilibrium binding constant (K_d_) determined from same dataset as (D), paired *t*-test. (**G**) Effect of MβCD (10 mM, 45 min) on exendin-4-induced cAMP production in stable HEK-SNAP-GLP-1R cells, 30 min stimulation, normalized to forskolin (FSK, 10 µM), *n*=4, E_max_ compared by paired *t*-test. (**H**) As for (G), but in INS-1 832/3 cells, 10 min stimulation in the presence of IBMX (500 µM), *n*=6, * E_max_ compared by paired *t*-test. (**I**) Confocal analysis of SNAP-GLP-1R internalization in MIN6B1 cells stably-expressing SNAP-GLP-1R labeled with SNAP-Surface 549 probe (red) for 30 min and then incubated with or without MβCD (10 mM, 1 hour) before stimulation with 100 nM exendin-4 for 15 min. Nuclei (DAPI), blue; size bars, 10 µm. (**J**) DERET measurements to quantify effect of MβCD (10 mM, 45 min) on exendin-4-induced (100 nM) SNAP-GLP-1R internalization in stable SNAP-GLP-1R-expressing HEK293 cells, expressed as HTRF ratio, *n*=4, 4-parameter logistic fit shown and used to quantify E_max_ and t_1/2_, compared by paired *t*-tests. (**L**) Confocal analysis of SNAP-GLP-1R internalization in wild-type (WT) and β-arrestin-less (βarr1/2 KO) HEK293 cells stably-expressing SNAP-GLP-1R labeled as in (I) and treated with or without MβCD before stimulation with 100 nM exendin-4 for 15 min. Nuclei (DAPI), blue; size bars, 10 µm. (**M**) TIRF microscopy analysis of plasma membranes from wild-type and β-arrestin-less HEK293 cells stably-expressing SNAP-GLP-1R and transiently-transfected with µ2-HA-WT, which codes for the µ2 domain of the clathrin adaptor AP2 fused to an HA-tag (AP2-HA), labeled with SNAP-Surface 488 (green) and stimulated for 1 min with 100 nM exendin-4 prior to fixation and HA-tag immunofluorescence (red); size bars, 10 µm. (**N**) Electron micrograph depicting a representative image of a clathrin-coated pit-localized gold-labeled SNAP-GLP-1R (arrow) in MIN6B1 cells stably-expressing SNAP-GLP-1R and stimulated for 1 min with 100 nM exendin-4); size bar, 100 nm. Unless stipulated, data are depicted as mean ± SEM; *p<0.05, ***p<0.001 by statistical test indicated in the text.

To investigate the effect of nanodomain recruitment on GLP-1R cellular behavior, we performed a series of assays in cells treated with or without methyl-β-cyclodextrin (MβCD), which disrupts the plasma membrane domain architecture via sequestration of cholesterol and other lipids (38). Efficiency of cholesterol depletion by MβCD was determined by filipin staining and quantified biochemically (Supplementary Figure 3A,B). MβCD treatment did not alter receptor surface expression levels (Supplementary Figure 3A,C).

**Figure 3.**
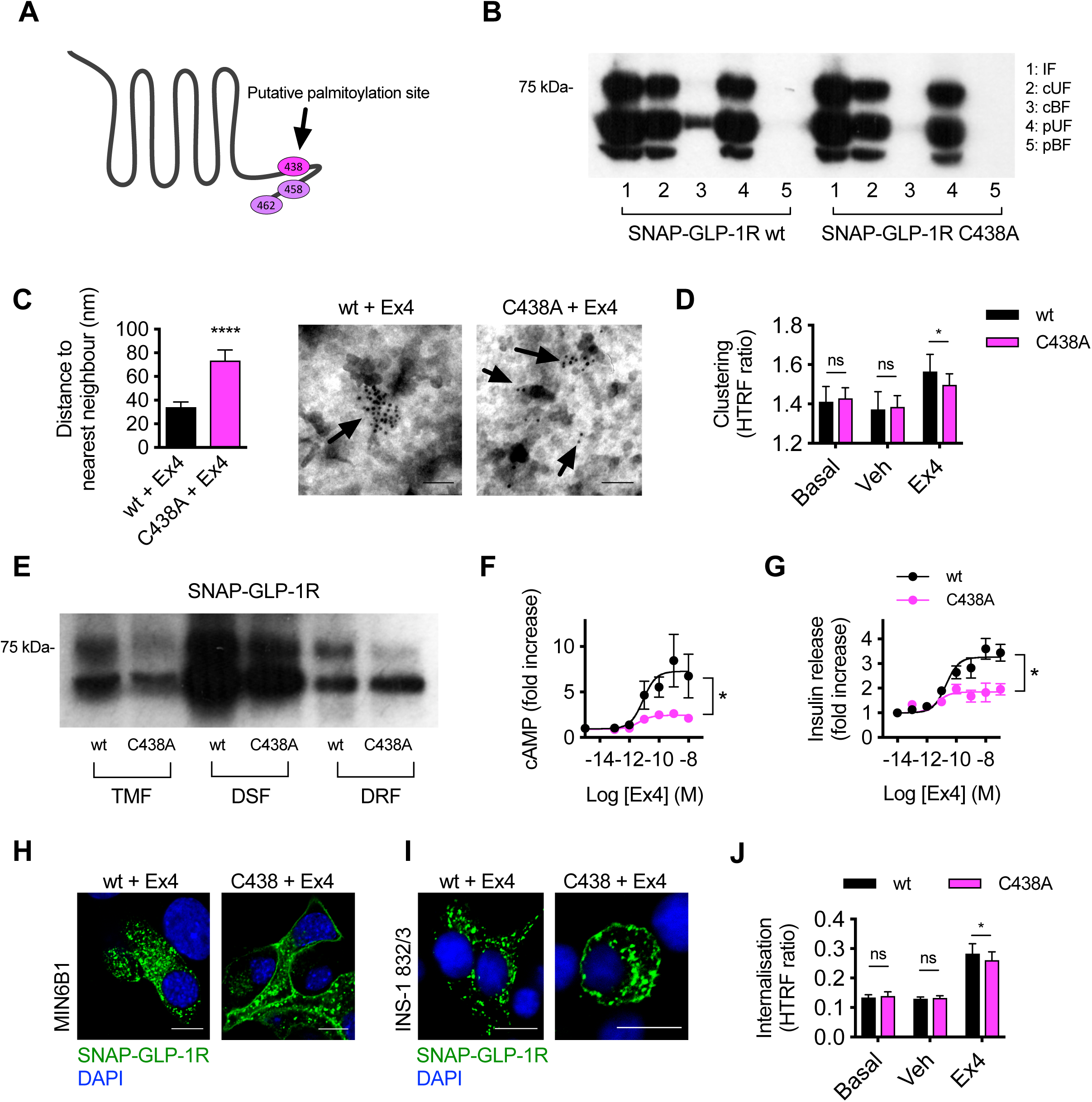
Effects of C438A palmitoylation-null mutant on clustering, membrane nanodomain recruitment, signaling and internalization properties of SNAP-GLP-1Rs. (**A**) Cartoon showing C-terminal cysteine residues in the GLP-1R cytoplasmic tail, with C438 highlighted as the putative palmitoylation site. (**B**) Analysis of SNAP-GLP-1R palmitoylation levels in CHO-K1 cells stably expressing SNAP-GLP-1R wild-type (wt) or C438A mutant and treated with 100 nM exendin-4 (Ex4) for 10 min. IF, input fractions; cUF, cleaved unbound fractions; cBF, cleaved bound fractions (corresponding to the palmitoylated pool); pUF, preserved unbound fractions; pBF, preserved bound fractions. (**C**) Distance to the nearest neighbor (left), and representative images of gold-labelled SNAP-GLP-1Rs (arrows, right), from EM analysis of 2-D plasma membrane sheets isolated from HEK293 cells stably-expressing SNAP-GLP-1R wild-type or C438A mutant after gold labeling of SNAP-tagged receptors and treatment with 100 nM exendin-4 for 2 min, data from a minimum of *n*=300 gold particles per condition, unpaired *t*-test. (**D**) Exendin-4 (100 nM) -induced clustering of wild-type or C438A SNAP-GLP-1Rs transiently-transfected in HEK293 cells, measured by HTRF, showing average HTRF ratios over a 10 min period pre-agonist addition (“Basal”) or over the last 10 min of a 30 min stimulation with vehicle or exendin-4, *n*=4, two-way repeat measures ANOVA with Sidak’s test. Kinetic data shown in Supplementary Figure 3D. (**E**) SNAP-GLP-1R wt *vs.* C438A distribution within total membrane (TMF), detergent-soluble (DSF) and detergent-resistant (DRF) fractions isolated from INS1 832/3 cells with endogenous GLP-1R expression deleted by CRISPR/Cas9 and stably-expressing either wild-type or C438A SNAP-GLP-1R and treated with 100 nM exendin-4 for 5 min. (**F**) Exendin-4-induced cAMP in INS-1 832/3 cells with endogenous GLP-1R expression deleted by CRISPR/Cas9 stably-expressing wild-type or C438A SNAP-GLP-1R, 10 min exendin-4 stimulation with 500 µM IBMX, *n*=4, 3-parameter fits shown, E_max_ compared by paired *t*-test. (**G**) Exendin-4-induced insulin secretion in INS-1 832/3 cells with endogenous GLP-1R expression deleted by CRISPR/Cas9 and stably expressing wild-type or C438A SNAP-GLP-1R; 16-hour stimulation at 11 mM glucose, expressed relative to 11 mM glucose alone, 3-parameter fits shown, E_max_ compared by paired *t*-test. (**H**, **I**) Confocal analysis of SNAP-GLP-1R wild-type *vs.* C438A internalization in MIN6B1 (**H**) and INS1 832/3 (**I**) cells with endogenous GLP-1R expression deleted by CRISPR/Cas9 and expressing wt or C438A mutant SNAP-GLP-1R following stimulation with 100 nM exendin-4 for 15 min. Nuclei (DAPI), blue; size bars, 10 µm. (**J**) Exendin-4 (100 nM) -induced internalization of wild-type or C438A SNAP-GLP-1R transiently transfected in HEK293 cells, measured by DERET, showing average HTRF ratios over a 10 min period pre-agonist addition (“Basal”) or over last 10 min of a 30 min stimulation with vehicle or exendin-4, *n*=7, two-way repeat measures ANOVA with Sidak’s test; kinetic data shown in Supplementary Figure 3M. Data are depicted as mean ± SEM; ns=non-significant, *p<0.05, ****p<0.0001, by statistical test indicated in the text.

MβCD treatment in HEK293 cells reduced both the magnitude and the rate of exendin-4-induced SNAP-GLP-1R clustering, as determined by HTRF (Figure 2A); a lower concentration of MβCD also slowed SNAP-GLP-1R clustering in CHO-K1 cells (Supplementary Figure 3D), albeit to a lesser extent. Moreover, MβCD treatment reduced both basal and exendin-4-induced NR12S-linked increases in TR-FRET from the liquid-ordered-associated portion of the spectrum (Figure 2B,C). As the presence of further binding sites in the immediate vicinity of a ligand that dissociates from its receptor could increase the chance of its “rebinding”, receptor clustering might increase the apparent agonist residence time (39). In keeping with this possibility, the apparent binding kinetics of fluorescently-labeled exendin-4-FITC (19) were notably altered by MβCD, with an increase in the agonist dissociation rate seen in both HEK293 (Figure 2D-F, TR-FRET traces shown in Supplementary Figure 3E) and CHO-K1 (Supplementary Figure 3F-I) cells stably expressing SNAP-GLP-1R, as determined by TR-FRET (40), resulting in an overall reduction in binding affinity.

We next measured the impact of MβCD treatment on GLP-1R-induced signaling responses. In HEK293 cells stably expressing SNAP-GLP-1R, and in the beta cell lines INS-1 832/3 and MIN6B1 with endogenously-expressed GLP-1R, acute cAMP responses to exendin-4 were reduced by MβCD (Figure 2G,H, Supplementary Figure 3J). Furthermore, AKAR4-Lyn and ^T^Epac^VV^ FRET biosensor measurements indicated that the impact of MβCD treatment on nanodomain-associated PKA activation was greater than the effect on total cAMP (39±6% *versus* 16±3% reduction in E_max_ after MβCD treatment for AKAR4-Lyn and ^T^Epac^VV^ responses, respectively) (Supplementary Figure 3K,L).

Membrane nanodomains have also been implicated in coupling receptor signaling with endocytosis (41). Confocal microscopy analysis of SNAP-GLP-1R stably expressed in MIN6B1 beta cells showed a significant reduction in exendin-4-induced SNAP-GLP-1R internalization after treatment with MβCD (Figure 2I). To quantify the effect of MβCD on SNAP-GLP-1R endocytosis kinetics, we used diffusion-enhanced resonance energy transfer (DERET), which measures disappearance from the cell surface of lanthanide-labeled SNAP-GLP-1Rs via reductions in energy transfer to extracellular fluorescein (25). This assay indicated a significant reduction in the SNAP-GLP-1R internalization rate in HEK293 cells with MβCD (Figure 2J), with a similar pattern seen in CHO-K1 cells (Supplementary Figure 3M,N).

Recruitment of β-arrestins to active GPCRs has traditionally been considered as a means of receptor desensitization coupled to clathrin-dependent endocytosis (37). In a previous study, we detected a very transient delay on GLP-1R internalization in cells lacking both β-arrestin-1 and -2, out of keeping with the more marked effects on signaling responses, which we now attribute primarily to avoidance of β-arrestin-mediated desensitization at the plasma membrane (19). In view of our latest observation of the profound effect of MβCD on GLP-1R internalization, we re-assessed the contribution of β-arrestins to GLP-1R internalization with and without cholesterol depletion by MβCD. Analysis of SNAP-GLP-1R localization by confocal microscopy in wild-type versus β-arrestin-1/2 knockout HEK293 cells (42) stably-expressing SNAP-GLP-1R showed at most very minor differences in SNAP-GLP-1R internalization propensity following stimulation with exendin-4. In contrast, this was substantially inhibited in both cell types after treatment with MβCD (Figure 2L), suggesting that the preservation of membrane nanodomain organization, rather than the recruitment of β-arrestins, is critical to sustain SNAP-GLP-1R endocytosis.

Endocytosis of receptors previously recruited to membrane nanodomains can occur via a range of clathrin-dependent and clathrin-independent pathways, with the specific pathway utilized in each case being particular to each individual receptor (43-46). β-arrestins classically couple GPCRs to clathrin via AP2 (47) so, in view of the persistence of GLP-1R endocytosis in the absence of β-arrestins, we aimed to investigate the route for GLP-1R endocytosis by performing TIRF microscopy analysis of SNAP-GLP-1R in HEK293 cells expressing SNAP-GLP-1R and HA-tagged AP2 following 1 minute stimulation with exendin-4. We found clustered SNAP-GLP-1Rs greatly co-localized with AP2-HA loci at the plasma membrane of both wild-type and β-arrestin-1/2 knockout cells (Figure 2M), indicating that the main pathway for entry of GLP-1Rs is indeed clathrin-mediated, but does not require β-arrestins. In order to confirm these results in beta cells, we performed EM analysis in sections from SNAP-GLP-1R-expressing MIN6B1 cells after gold labeling as in Figure 1. Stimulation with exendin-4 for 1 minute resulted in localization of gold particles to clathrin-coated pits (Figure 2N).

In order to further analyze the specific role of GLP-1R palmitoylation on exendin-4-triggered cellular effects, we generated a mutant version of SNAP-GLP-1R by site-directed mutagenesis to replace the cysteine 438 at the C-terminus of the receptor, previously described as its only palmitoylation site (20), with alanine (C438A; Figure 3A). We first confirmed that mutant SNAP-GLP-1R C438A is no longer palmitoylated following exendin-4 stimulation (Figure 3B). Note that expression levels of both receptor types were similar, either after transient transfection, or in the stable cell lines used in this study (Supplementary Figure 4A-D), suggesting that constitutive palmitoylation is not a requirement for the efficient delivery of the GLP-1R to the plasma membrane, unlike for some other GPCRs (48). Additionally, we did not detect any significant differences in binding affinity to exendin-4 with SNAP-GLP-1R C438A mutant *versus* wild-type receptors (Supplementary Figure 4E-G).

**Figure 4.**
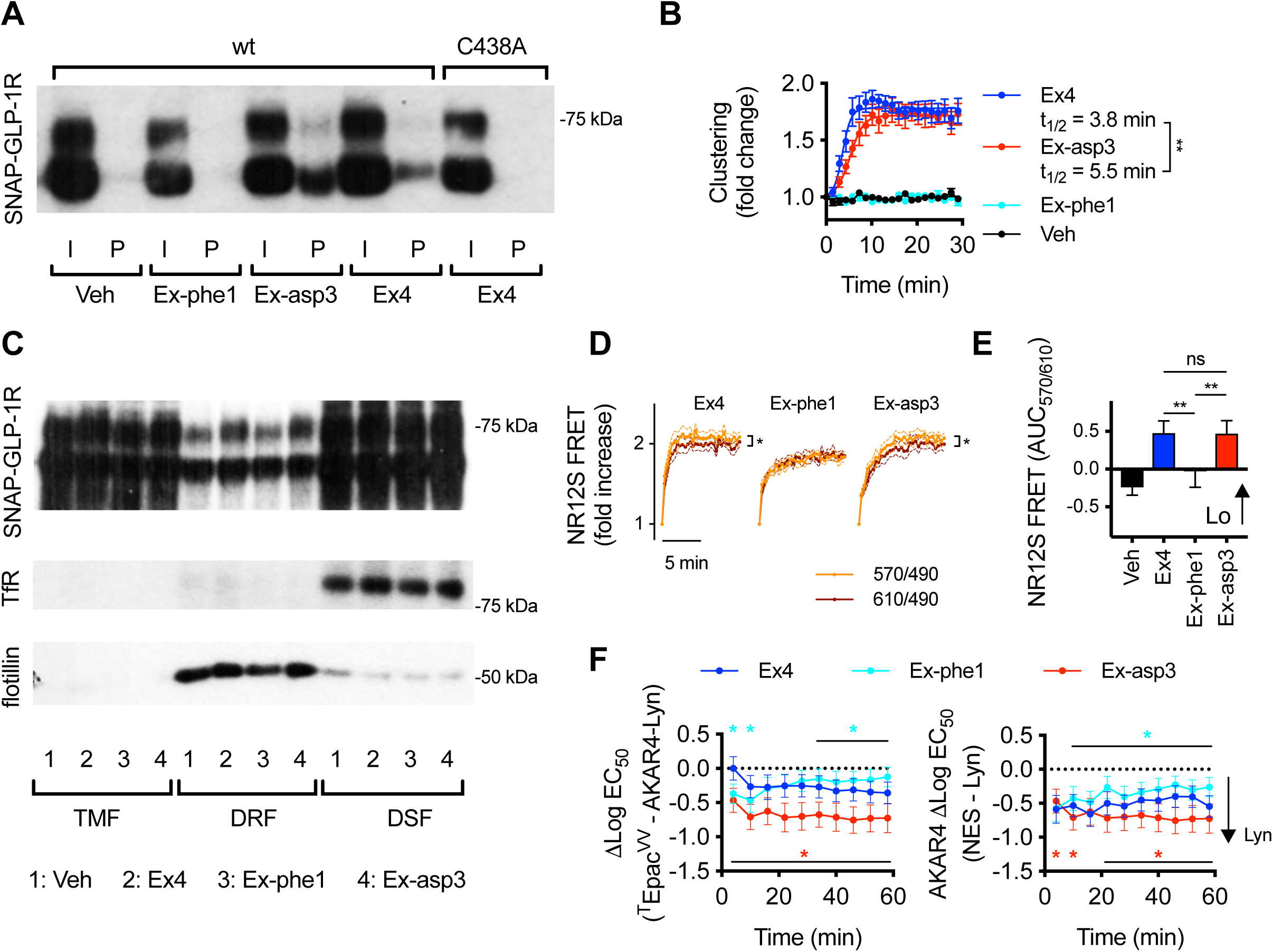
Control of SNAP-GLP-1R palmitoylation, clustering and membrane nanodomain recruitment and signaling by biased exendin-4 agonists. (**A**) Total input (I) and palmitoylated (P) SNAP-GLP-1R fractions from CHO-K1 cells stably-expressing SNAP-GLP-1R wild-type (wt) or C438A and treated with vehicle (Veh) or 100 nM of exendin-4 (Ex4), exendin-phe1 (Ex-phe1) or exendin-asp3 (Ex-asp3) for 10 min. (**B**) HTRF clustering measurement of clonal CHO-SNAP-GLP-1R cells treated with the indicated agonist (100 nM), expressed relative to individual well baseline, *n*=6, 4-parameter fit used to quantify t_1/2_, compared by paired *t*-test. (**C**) SNAP-GLP-1R distribution within total membrane (TMF), detergent-resistant (DRF) and detergent-soluble (DSF) fractions isolated from MIN6B1 cells stably-expressing SNAP-GLP-1R treated with vehicle or 100 nM of the indicated agonist for 2 min, with transferrin receptor (TfR) as a marker of membrane non-raft enrichment and flotillin as a marker of membrane raft enrichment. Quantification is shown in Supplementary Figure 4A. (**D**) Kinetic traces of NR12S FRET in stable HEK-SNAP-GLP-1R cells treated with the indicated agonist (100 nM), expressed as ratio of signals at 570 nm to 490 nm (representing Lo signal), or at 570 nm to 490 nm (representing Ld signal), and normalized to individual well baseline, *n*=6, paired *t*-tests comparing plateau determined from one-phase association fits. (**E**) Alternative analysis of data shown in (D), with AUC calculated from ratio of 570 and 610 signals, randomized block one-way ANOVA with Tukey’s test. (**F**) Summary of relative potency changes in CHO-SNAP-GLP-1R cells treated with indicated agonist with ^T^Epac^VV^ *vs.* AKAR4-Lyn responses, or AKAR4-NES *vs.* AKAR4-Lyn responses, determined by subtracting Log EC_50_ for each biosensor at each time-point, *n*=5, two-way repeat measures ANOVA with Dunnett’s test *vs.* exendin-4. Data are depicted as mean ± SEM; ns=non-significant, *p<0.05, **p<0.01, by statistical test indicated in the text.

In keeping with a putative role for palmitoylation in contributing to the formation of receptor clusters, EM analysis of nearest neighbors in 2-D plasma membrane sheets from gold-labeled HEK293 cells stably expressing each SNAP-tagged receptor showed a reduced clustering effect of exendin-4 with the C438A mutant (Figure 3C). This observation was supported by HTRF measurements, which showed that receptor clustering in HEK293 cells transiently expressing wild-type or C438A SNAP-GLP-1R was similar under basal or vehicle-treated conditions, but the expected exendin-4-induced increase was less marked with the C438A mutant (Figure 3D, Supplementary Figure 4H). Note that, in the latter assay, agonist-induced changes in HTRF ratios were small in comparison to those seen with stably-expressing cells (e.g. in Figure 2A), and therefore full kinetic analyses could not be performed reliably; statistical comparisons were consequently made with average signals detected over the final 10 minutes of a 30-minute treatment period. Concomitantly, recruitment of SNAP-GLP-1R C438A to detergent-resistant fractions following exendin-4 stimulation was reduced compared to SNAP-tagged wild-type receptor when expressed in INS-1 832/3 cells engineered by CRISPR/Cas9 to delete endogenous GLP-1R (49), thereby avoiding confounding co-expression of two forms of the receptor (Figure 3E). Similar findings were made using CHO-K1 cells stably expressing C438A or wild-type SNAP-GLP-1Rs (Supplementary Figure 4I).

We then determined the impact of the C438A mutation on SNAP-GLP-1R signaling, which has variously been reported to result in a reduction in cAMP signaling (20, 50) or to have no effect (51). In accordance with the former reports, and consistent with the effect of cholesterol depletion that we identified in Figure 2, we found that cAMP production was reduced with the C438A palmitoylation mutant compared to wild-type SNAP-GLP-1R in INS-1 832/3 cells lacking endogenous GLP-1R (Figure 3F). Critically, this signaling defect in beta cells translated into significant reductions in exendin-4-induced insulin secretion (Figure 3G), indicating that GLP-1R palmitoylation may play an important role in physiological glucose homeostasis or response to pharmacological GLP-1R agonists in diabetes. Consistently, recruitment of β-arrestin-2 to SNAP-GLP-1R C438A was reduced compared to wild-type, as measured in exendin-4-stimulated HEK293 cells stably expressing β-arrestin-2-EA and transiently transfected with ProLink-tagged wild-type or C438A SNAP-GLP-1R (Supplementary Figure 4J). Total cAMP production was also reduced in CHO-K1 cells with stable expression of C438A SNAP-GLP-1R compared to wild-type when measured biochemically (Supplementary Figure 4K); we also used the ^T^Epac^VV^ and AKAR4-Lyn FRET biosensors to measure localized and total responses in these cells, which indicated that the signaling defect particularly affected nanodomain-associated PKA activation (Supplementary Figure 4L-N), although the small signal changes in this assay precluded any kinetic comparisons, and dose-response curves could only be constructed from averaged response data over the entire 30-minute incubation period.

Finally, we compared the trafficking properties of wild-type *versus* C438A mutant SNAP-GLP-1R when expressed in MIN6B1 cells in which we knocked out endogenous GLP-1R by CRISPR/Cas9 (Supplementary Figure 4O), as well as in the endogenous GLP-1R-null INS-1 832/3 cells described earlier. We detected a small and transient delay in the internalization of the C438A palmitoylation mutant in both beta cell lines (Figure 3H,I), with no effect upon receptor recycling following internalization (Supplementary Figure 4P,Q). Internalization measurements by DERET of wild-type or C438A SNAP-GLP-1Rs again revealed subtly reduced internalization propensity with the palmitoylation-deficient mutant in HEK293 cells (Fig. 3J, Supplementary Figure 4R). A similar minor delay in internalization was seen with CHO-K1 cells stably expressing SNAP-GLP-1R wild-type *versus* C438A (Supplementary Fig 4S,T).

We have previously investigated the effects of a panel of exendin-4-based biased GLP-1R agonists with marked differences in binding kinetics, preference for cAMP *versus* β-arrestin recruitment, receptor internalization and recycling (19). Although they were measured acutely, the bias profiles were predictive of the ability of each agonist to support long-term GLP-1R signaling responses and insulin secretion over many hours from beta cells, with lower affinity, slow-internalizing agonists more efficacious for longer-term responses despite being less potent acutely. Given that differential palmitoylation and/or nanodomain partitioning has been linked to the establishment of bias (22), and prompted by the apparent similarities between the trafficking and signaling effects of GLP-1R biased agonists and those described here following disruption of membrane nanodomains or GLP-1R palmitoylation, we next explored the possibility that two exemplar agonists from this panel, exendin-phe1 and exendin-asp3, selected as displaying opposite bias effects, might influence GLP-1R palmitoylation, clustering and/or recruitment to membrane nanodomains compared to the reference agonist exendin-4.

We first analyzed the level of SNAP-GLP-1R palmitoylation elicited by these three agonists in CHO-K1 cells, and found that stimulation with exendin-asp3, a high affinity agonist biased towards β-arrestin recruitment and propensity for internalization, increased SNAP-GLP-1R palmitoylation compared to exendin-4, while stimulation with exendin-phe1, a low affinity agonist biased in the opposite direction, had a reduced effect (Figure 4A). Broadly in keeping with this observation, SNAP-GLP-1R clustering was rapid with both exendin-4 and exendin-asp3, but virtually absent with exendin-phe1 (Figure 4B).

We next examined whether a similar pattern was present for membrane raft partitioning by analysis of SNAP-GLP-1R levels in detergent-resistant *versus* detergent-soluble membrane fractions from MIN6B1 cells stably expressing SNAP-GLP-1R. We found that a larger proportion of SNAP-GLP-1R segregated to detergent-resistant fractions following stimulation with exendin-asp3 and exendin-4 compared to exendin-phe1 (Figure 4C, quantified in Supplementary Figure 5A), underlying the capacity of the biased agonists to control both the level of palmitoylation and nanodomain recruitment of the receptor. In keeping with this biochemical assay, kinetic monitoring of NR12S FRET in HEK293 cells expressing SNAP-GLP-1R treated with biased agonists indicated a small but consistently greater increase in signal from liquid-ordered (570 nm) *versus* liquid-disordered (610 nm) spectral regions with exendin-4 and exendin-asp3, in comparison to exendin-phe1 (Figure 4D,E).

**Figure 5.**
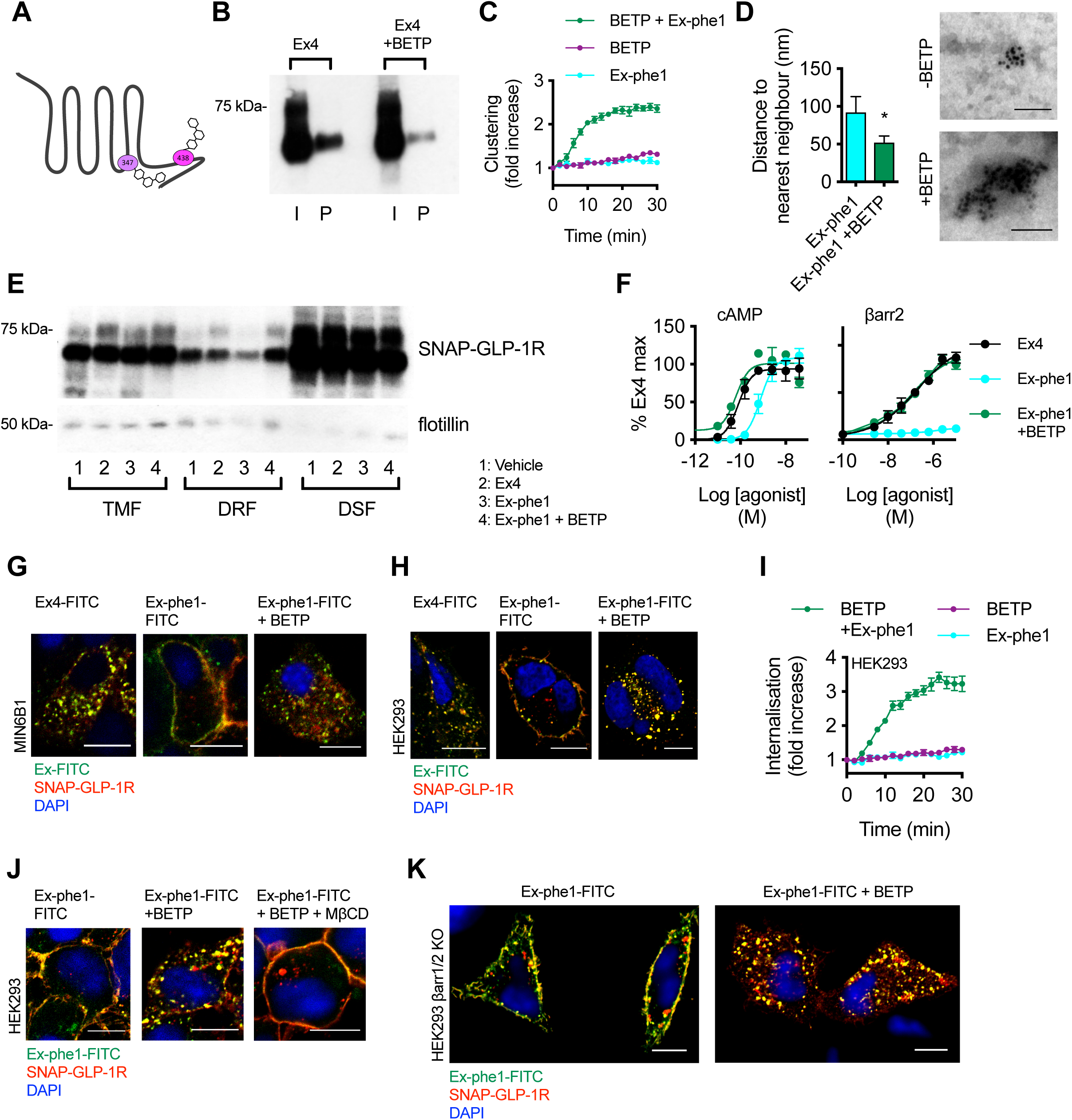
Effect of BETP on SNAP-GLP-1R clustering, nanodomain recruitment, signaling and internalization responses. (**A**) Cartoon depicting sites of covalent modification of GLP-1R by BETP at C347 and C438, as per (50). (**B**) Total input (I) and palmitoylated (P) SNAP-GLP-1R fractions from CHO-K1 cells stably expressing SNAP-GLP-1R treated with 100 nM exendin-4 (Ex4) for 10 min in the presence or absence of 10 µM BETP. (**C**) SNAP-GLP-1R clustering measured by HTRF in monoclonal CHO-SNAP-GLP-1R cells treated with exendin-phe1 (Ex-phe1, 100 nM), BETP (10 µM), or both, expressed ratiometrically and after normalization to individual well baseline, *n*=4. (**D**) Distance to the nearest neighbor (left), and representative images of gold-labelled SNAP-GLP-1Rs (right), from EM analysis of 2-D plasma membrane sheets isolated from MIN6B1 cells stably-expressing SNAP-GLP-1R following gold labeling of SNAP-tagged receptors and treatment with 100 nM exendin-phe1 with and without BETP (10 µM) for 2 min, data is from a minimum of *n*=155 (-BETP) or *n*=330 (+ BETP) gold particles per condition, unpaired *t*-test. (**E**) SNAP-GLP-1R distribution within total membrane (TMF), detergent-resistant (DRF) and detergent-soluble (DSF) fractions isolated from MIN6B1 cells stably expressing SNAP-GLP-1R treated with vehicle (Veh) or 100 nM of the indicated agonist with and without BETP (10 µM) for 2 min, with flotillin as a marker of membrane raft enrichment. (**F**) Effect of BETP (10 µM) on exendin-phe1-induced cAMP and β-arrestin-2 recruitment in PathHunter GLP-1R cells, 30 min incubation, 4-parameter fit shown, *n*=3. (**G**, **H**) Confocal analysis of indicated FITC-agonist (green) and SNAP-GLP-1R (red) internalization in MIN6B1 (**G**) and HEK293 (**H**) cells stably expressing SNAP-GLP-1R following stimulation with 100 nM of the indicated FITC-agonist with or without BETP (10 µM) for 10 min (**G**) or 40 min (**H**). Nuclei (DAPI), blue; size bars, 10 µm. (**I**) SNAP-GLP-1R internalization measured by DERET in stable HEK-SNAP-GLP-1R cells treated with exendin-phe1 (100 nM), BETP (10 µM), or both, expressed ratiometrically and after normalization to individual well baseline, *n*=4. (**J**) Confocal analysis of exendin-phe1-FITC (green) and SNAP-GLP-1R (red) internalization in HEK293 cells stably expressing SNAP-GLP-1R following stimulation with 100 nM of exendin-phe1-FITC for 10 min with or without BETP (10 µM) and MβCD (10 mM, 1-hour pre-incubation). Nuclei (DAPI), blue; size bars, 10 µm. (**K**) Confocal analysis of exendin-phe1-FITC (green) and SNAP-GLP-1R (red) internalization in HEK293 β-arrestin-less cells stably expressing SNAP-GLP-1R following stimulation with 100 nM of exendin-phe1-FITC for 40 min with or without BETP (10 µM). Nuclei (DAPI), blue; size bars, 10 µm. Data are depicted as mean ± SEM; *p<0.05, by statistical test indicated in the text.

Responses to each biased agonist at multiple doses were next measured in real-time in CHO-K1 cells expressing SNAP-GLP-1R and the biosensors ^T^Epac^VV^, AKAR4-NES, or AKAR4-Lyn (Supplementary Figure 5B), allowing the construction of sequential dose-response curves across the entire stimulation period, in order to observe changes in potency over time (Supplementary Figure 5C). Readings were taken every two minutes, and signals were combined into six-minute bins to improve precision. Using these data, we assessed whether the differential membrane raft translocation of biased agonists could control spatial organization of cAMP signaling. Bias towards nanodomain-specific AKAR4-Lyn signaling, compared to both ^T^Epac^VV^ and AKAR4-NES, was indeed greatest with exendin-asp3, and least with exendin-phe1, particularly at later time points (Figure 4F).

An apparent link between GLP-1R binding kinetics and agonist-induced translocation of activated receptors to cholesterol-rich nanodomains could be inferred from the similar behaviors of the GLP-1R when activated either by 1) the low affinity agonist exendin-phe1, or 2) exendin-4 after pre-treatment with affinity-reducing MβCD. Prompted by this, as well as by our previous observation of the capacity of the GLP-1R-specific PAM BETP to increase the residence time of exendin-4 to the GLP-1R (19), we next investigated whether BETP would be able to reverse the reduced detergent-resistant fraction recruitment and internalization phenotype of exendin-phe1-stimulated SNAP-GLP-1R. Fortuitously, BETP also presents an opportunity to independently examine the role of GLP-1R palmitoylation on downstream events, as it crosses the plasma membrane to covalently bind cysteine 438 (i.e. occupying the putative GLP-1R palmitoylation site) whilst exerting its allosteric effect via binding to cysteine 347 (50) (see Figure 5A for a schematic of BETP binding sites at the GLP-1R). Indeed, as expected from its known targeting of cysteine 438, BETP caused a significant reduction in exendin-4-induced palmitoylation (Figure 5B), indicating that it is indeed able to block transfer of palmitate to the GLP-1R by binding to its known palmitoylation site.

We next aimed to assess whether binding of BETP to the GLP-1R could “rescue” some of the pharmacological effects of exendin-phe1. In keeping with results from CHO-K1 cells expressing SNAP-GLP-1R (Figure 4B), exendin-phe1 failed to induce significant GLP-1R clustering in HEK293 cells stably expressing SNAP-GLP-1R (Figure 5C). The effect of BETP alone was marginal; however, co-application of exendin-phe1 with BETP resulted in rapid SNAP-GLP-1R clustering, comparable to that obtained with exendin-4 (e.g. see Figure 2A). These results were confirmed by EM analysis of gold-labeled 2-D membrane sheets from MIN6B1 cells stably expressing SNAP-GLP-1R and stimulated with exendin-phe1 in the presence or absence of BETP (Figure 5D). We have previously noted how BETP slows down the dissociation of orthosteric GLP-1R agonists from the receptor (19, 23), and in the present study we have found that the dissociation rate of exendin-phe1-FITC from SNAP-GLP-1R is indeed notably slower with BETP treatment, which is partly mitigated by an apparent concomitant slower association rate, resulting in a non-significant increase to overall binding affinity (Supplementary Figure 6A-D). Whilst the mechanism by which this occurs is not currently established, the observed increase in BETP-mediated GLP-1R clustering might contribute to agonist rebinding, as highlighted earlier from the contrary effect of MβCD (Figure 2D-F).

We next examined whether BETP was able to rescue the reduced SNAP-GLP-1R segregation to membrane nanodomains elicited by exendin-phe1 (Figure 5E). In agreement with the results obtained in Figure 4C-E, stimulation with exendin-phe1 resulted in reduced recruitment of SNAP-GLP-1R to detergent-resistant fractions of MIN6B1 cells stably-expressing SNAP-GLP-1R compared to exendin-4; however, addition of BETP was able to reverse the exendin-phe1 phenotype, with SNAP-GLP-1Rs now being recruited to membrane rafts to the same extent as with exendin-4. BETP was also able to trigger a marked increase in exendin-phe1-induced cAMP and β-arrestin-2 recruitment to the GLP-1R, with the resulting responses again comparable to those of exendin-4 (Figure 5F).

Stimulation of CHO-K1 cells expressing SNAP-GLP-1R with increasing concentrations of BETP alone (without any orthosteric agonist) resulted in activation of both cytoplasmic (AKAR4-NES) and raft-associated (AKAR4-Lyn) PKA; however, the lower threshold for activation of AKAR4-Lyn suggested that the effect was relatively nanodomain-specific (Supplementary Figure 6E,F). Additionally, in allosteric mode at a lower concentration, BETP modestly but selectively potentiated the AKAR4-Lyn response to exendin-phe1, without affecting that of AKAR4-NES (Supplementary Figure 6G,H).

Finally, we examined whether BETP would be able to reverse the reduced internalization previously observed with exendin-phe1-stimulated SNAP-GLP-1R (19). Here we used FITC-conjugates of both exendin-4 and exendin-phe1, in conjunction with surface-labeled SNAP-GLP-1R, in order to simultaneously monitor the endocytic behaviors of both agonist and receptor. As expected, stimulation with exendin-phe1-FITC triggered significantly less receptor internalization than with exendin-4-FITC in MIN6B1 cells stably expressing SNAP-GLP-1R, but addition of BETP restored an exendin-4-like rapid internalization phenotype to exendin-phe1-FITC (Figure 5G). BETP alone (without orthosteric agonist) had no effect on GLP-1R internalization (Supplementary Figure 7A). Congruent results were obtained using HEK293 (Figure 5H,I) and CHO-K1 (Supplementary Figure 7B) cells stably expressing SNAP-GLP-1R.

Interestingly, BETP remained capable of stimulating internalization of the C438A SNAP-GLP-1R mutant in both stably-expressing CHO-K1 cells and transiently-transfected HEK293 cells (Supplementary Figure 7C,D). This finding is in keeping with the current understanding that the pharmacology of BETP is mediated via binding to cysteine 347 rather than 438 (50), and further suggests that C438 palmitoylation (reduced with BETP as per Figure 5B) is not an obligatory event for GLP-1R endocytosis. However, the BETP-induced internalization of exendin-phe1-stimulated SNAP-GLP-1Rs still relied on cholesterol-dependent membrane compartmentalization, as pre-treatment with MβCD inhibited this process to the same extent as for exendin-4 (Figure 5J), and was fully recapitulated in HEK293 cells even in the absence of β-arrestins (Figure 5K). As for exendin-4 (Figure 2M,N), SNAP-GLP-1R internalization induced by exendin-phe1 in the presence of BETP primarily used a clathrin-dependent pathway of endocytosis, as it involved engagement of the clathrin adaptor AP2, seen by TIRF microscopy analysis in MIN6B1 cells expressing SNAP-GLP-1R (Supplementary Figure 7E), and resulted in localization of gold-labeled SNAP-GLP-1Rs to clathrin-coated pits, seen by EM analysis in SNAP-GLP-1R-expressing HEK293 cells (Supplementary Figure 7F).

## Discussion

Class B GPCRs such as the GLP-1, glucagon and GIP receptors play crucial roles in the control of glucose and energy metabolism, and are key investigational targets for the treatment of several metabolic disorders including insulin resistance, obesity and type 2 diabetes (52). Of these, the GLP-1R is the only one for which approved therapies are in current clinical use (53). With the current sharp rise in the worldwide incidence of both type 2 diabetes and obesity (54), there is a pressing need to develop more effective drugs with fewer associated side effects. The discovery of biased signaling, by which a GPCR can transduce specific signaling profiles by favoring one pathway or behavior over another (55), opens up new avenues to achieve deeper control of GLP-1R action and cellular responses, as previously shown by us (19), and others (56).

In this context, the role of the plasma membrane lipid environment in the control of GPCR functional selectivity remains a relatively underexplored area, due primarily to the technical challenges associated with the study of lipid-protein cross-talk (7). Despite this, emerging evidence using newly developed methods places lipid nanodomains as key regulators of GPCR signaling (2, 5, 57-59).

In this study, we have unveiled that the palmitoyl acylation of GLP-1R at cysteine 438 of its cytoplasmic tail, as well as its clustering in membrane rafts, is greatly potentiated by agonist binding, indicating that the GLP-1R interaction with the surrounding lipid environment is conditioned by its activation state. Interestingly, the same was not found with the closely related GIPR, which has a higher constitutive activity in the absence of agonist stimulation (60), potentially explaining the greater level of basal palmitoylation without further increase when stimulated with GIP. Palmitoylation is the only reversibly regulated post-translational lipid modification, and as such, it is widely considered as a dynamic raft-targeting mechanism (17, 61). Consequently, we observe a correlation between the level of agonist-induced GLP-1R palmitoylation and clustering within liquid-ordered nanodomains. In living cells, raft assemblies can be stabilized by specific oligomerization/clustering of proteins with little energy input, with protein raft affinity further enhanced by oligomerization (9), so that the processes of liquid-ordered nanodomain formation and receptor clustering might be mutually augmented. It is therefore possible that the agonist binding-triggered rise in GLP-1R clustering, in combination with the increased palmitoylation state of the agonist-bound receptor, heralds its recruitment to membrane rafts, which might explain the residual exendin-4-induced clustering observed after MβCD-dependent disruption of the membrane nanodomain architecture.

Membrane rafts are thought to compartmentalize cellular processes by contributing to the organization of signaling molecules. The necessity for the accumulation of certain downstream signaling effectors for efficient signal transduction might define raft-specific responses, with small variations often resulting in large changes at the cellular level (62). Additionally, receptors are known to undergo specific conformational changes upon interaction with lipid nanodomains. For example, binding to a specific raft lipid has been shown to allosterically change the conformation of the human EGF receptor (63). Thus, functional association of the GLP-1R with membrane rafts might potentially induce specific conformational changes that could modulate its function. In this context, our observation that exendin-4-induced cAMP signaling, as well as the main GLP-1R signal transducer, GαS, are both predominantly raft-associated, underlines the importance of agonist-induced GLP-1R nanodomain segregation in the optimization of GLP-1R signaling, with disturbances in this process predicted to carry deep consequences for the control of insulin secretion, as illustrated in the present study by the acute signaling and insulin secretion defects harbored by the C438A palmitoylation-null mutant.

One further way in which receptor clustering might influence signal generation relates to the potential modulation of ongoing agonist binding events. Firstly, crosstalk within oligomeric complexes might alter the binding properties of individual receptor protomers (64), although this typically manifests as negative cooperativity, i.e. with secondary protomers displaying reduced affinity, as previously described for the GLP-1R when in constitutive homodimeric configuration at the plasma membrane (29). Alternatively, aggregation of receptors within specific membrane regions could shift the behavior of a dissociating agonist towards rebinding to a nearby available receptor, rather than diffusing away into the extracellular space. This phenomenon could increase the apparent “residence time” of ligands at their receptor and prolong drug action (65). Here we provide some evidence to support the latter phenomenon, as disruption of GLP-1R clustering and membrane nanodomain segregation by MβCD leads to an apparent increase in the dissociation constant of exendin-4 to the GLP-1R. Conversely, allosteric augmentation of agonist residence time with the GLP-1R-specific PAM BETP is in itself capable of inducing nanodomain clustering, resulting in enhanced signaling from membrane rafts and potentiation of GLP-1R internalization, indicating that, as for the processes of receptor clustering and membrane raft segregation, these two phenomena are mutually regulated.

Modulation of GPCR post-translational modifications, as well as receptor clustering and recruitment to specific plasma membrane nanodomains, have previously been suggested as means by which biased signaling can be regulated (22). We found marked differences in GLP-1R clustering, nanodomain recruitment and internalization between biased exendin-4 agonists previously shown by us to exhibit significant differences in receptor binding affinity and signaling preferences (19), as well as distinct GLP-1R palmitoylation profiles, as shown in the present study. These effects were recapitulated to a certain extent by the palmitoylation-null C438A GLP-1R mutant. However, the observed reductions in receptor clustering, membrane raft segregation and signaling, as well as the reduction in β-arrestin recruitment and GLP-1R internalization, were noticeably milder with the latter compared to the biased agonist exendin-phe1. Of note, we did not detect any significant differences in exendin-4 binding kinetics with the GLP-1R C438A mutant *versus* the wild-type receptor, while exendin-phe1 displays a marked reduction in GLP-1R binding affinity compared to exendin-4 (19). It is important to point out that despite the role of GLP-1R palmitoylation in favoring receptor clustering in membrane rafts, this post-translational modification alone is not sufficient for membrane raft association. There are many examples of palmitoylated proteins that are not raft-associated (66), including the commonly used non-raft marker transferrin receptor. Moreover, as shown here, BETP is capable of inducing nanodomain clustering and enhancing raft signaling while at the same time resulting in the inhibition of GLP-1R palmitoylation, the latter presumably due to its covalent binding to the GLP-1R cysteine 438, among others (50), which would block any post-translational modification at this residue. Whether binding of BETP at cysteine 438 could functionally substitute for the absence of palmitic acid at this residue to promote nanodomain translocation was not formally assessed in the present study, but this appears improbable as its effect on GLP-1R endocytosis was preserved in the GLP-1R C438A mutant, which was also previously reported to show normal potentiation of partial agonist signaling responses in contrast to C347A (50). Taken together, the above considerations suggest that the control of GLP-1R recruitment to membrane nanodomains and its associated effects on receptor signaling are likely to encompass contributions from various factors. While we have not established the exact mechanism by which the exendin-4-based biased agonists influence the level of GLP-1R raft recruitment and clustering observed in this study, it is likely that a combination of factors, such as specific agonist-bound GLP-1R conformations that favor or stabilize receptor palmitoylation, differences in receptor occupancy related to binding affinity, differential engagement of activated receptors with spatially constrained G proteins (67), and potential agonist-specific differences in cytoskeletal rearrangements (68), underlie their different effects.

Nanodomain segregation is intimately linked with the control of receptor endocytosis (45). Accordingly, we found profound effects on agonist-induced GLP-1R internalization following cholesterol extraction with MβCD. Moreover, the GLP-1R internalization profiles exhibited by the biased exendin-4 agonists (19) closely correlate with their effects on GLP-1R membrane raft segregation and clustering, and the reduced GLP-1R internalization elicited by exendin-phe1 is fully rescued by the restoration of GLP-1R clustering and nanodomain segregation by BETP in a manner that still relies on the cholesterol-dependent preservation of membrane nanodomain architecture. Interestingly, the GLP-1R appears to follow a clathrin-dependent pathway of internalization for all the conditions tested. This is despite the fact that, while the recruitment of β-arrestins to the GLP-1R occurred preferentially from membrane nanodomains, indicative of a higher level of receptor activation within these raft regions and presumably associated with the β-arrestin-dependent GLP-1R desensitization previously observed by us (19), β-arrestins appeared to play a very minor role in GLP-1R endocytosis. It is worth noting that the GLP-1R harbors an AP2-binding domain in its C-terminal tail, and can interact directly with this clathrin adaptor, reducing the need for additional intermediaries (69). It remains to be established whether the receptor internalizes from cholesterol-rich membrane nanodomains by an atypical clathrin-mediated mechanism (44), or alternatively, as seen before for other GPCRs (70), its association with membrane rafts precedes its recruitment to clathrin-coated pits. Future single-particle tracking experiments to assess the GLP-1R dynamics at the plasma membrane of living cells (5) will be required to elucidate this.

The reduction in acute signaling seen with both the GLP-1R C438A palmitoylation-null mutant and the biased agonist exendin-phe1 can be easily reconciled with their failure to sustain optimal GLP-1R clustering and raft recruitment, which would preclude a rapid and synchronized signaling response. However, exendin-phe1 is also counter intuitively associated with long-term preservation of signaling and beneficial glucose lowering effects *in vivo* (19), whereas the insulin secretory capacity of the C438A mutant was markedly lower than wild-type when maximally stimulated with exendin-4 over several hours. Noting the more marked effects of exendin-phe1 *versus* the C438A mutant on GLP-1R β-arrestin recruitment, endocytosis and recycling (19), it would seem plausible that these are large enough in the case of exendin-phe1 to offset any negative effects on rapid acute signaling via a better preservation of a sizeable pool of re-sensitized surface receptors that do not undergo lysosomal degradation, and therefore allow for long-term potentiation of insulin secretion. Further experimental differences may also contribute, for example, over-expression of wild-type or C438A SNAP-GLP-1Rs in the INS-1 832/3 GLP-1R CRISPR/Cas9 model might not fully recapitulate endogenous expression of GLP-1R in the parental cells in which biased agonist testing was originally performed (19).

The results from the present study suggest that the cellular lipid composition plays important roles on the preservation of GLP-1R signaling. Lipotoxic insults, in particular exposure to palmitate, have previously been shown to result in disturbances in GLP-1-triggered effects on insulin secretion (71). Excess palmitate may not only disturb the distribution and partitioning of lipids at the plasma membrane, but also appears to induce hyper-palmitoylation of the GLP-1R in basal conditions within beta cells (see Supplementary Figure 1D), potentially resulting in increased GLP-1R desensitization that might contribute to the previously observed beta cell incretin resistance phenotype.

Additionally, it is important to note that raft size and composition depends on the specific membrane environment of each individual cell type. Investigating the possible role of the lipid environment on the regulation of GLP-1R function in the various cell types where it is expressed, such as hypothalamic neurons *versus* pancreatic beta cells, will be paramount to explain possible differences in GLP-1R behavior between these different cellular contexts.

In conclusion, in the present study we have employed biochemical assays in conjunction with high-resolution microscopy and spatially localized FRET biosensor experiments to establish the existence of an agonist binding-triggered GLP-1R palmitoylation and clustering mechanism leading to GLP-1R segregation to and signaling from plasma membrane nanodomains, and coupled to cholesterol-dependent GLP-1R internalization and desensitization. We have also established that this mechanism can be controlled by the modulation of agonist binding affinities, either with biased GLP-1R agonists or with the PAM BETP. Future investigations, including the identification of the specific palmitoyltransferase(s) and palmitoyl-protein thioesterase(s) responsible for the regulation of the agonist-induced GLP-1R palmitoylation in beta cells and other relevant cell types, as well as the use of model systems in order to fine-tune the plasma membrane lipid composition and nanodomain architecture, will be relevant to further develop our current understanding of the impact of the lipid composition on the spatiotemporal organization of GLP-1R signaling.

## Materials and Methods

### Agonists

Exendin-4 and exendin(9-39) were from Bachem, and custom peptides were from Insight Biotechnology. FITC-conjugates of exendin-4 and exendin-phe1 were coupled via K12, as previously described (19). BETP was from Sigma Aldrich.

### Cell culture and generation of stable cell lines

HEK293 cells (ECACC) were maintained in DMEM supplemented with 10% fetal bovine serum (FBS) and 1% penicillin/streptomycin. CHO-K1 cells (ECACC) were maintained in DMEM supplemented with 10% FBS, 20 mM HEPES, 1% non-essential amino acids, and 1% penicillin/streptomycin. PathHunter CHO-GLP-1R β-arrestin-2 reporter cells (DiscoverX) were maintained in the manufacturer’s Culture Medium 6. HEK293 β-arrestin-2-EA cells were generated by infection of HEK293 cells with viral retroparticles expressing β-arrestin-2 fused to EA (DiscoverX) and selected in hygromycin. Parental INS-1 832/3 cells, and INS-1 832/3 cells with endogenous GLP-1R or GIPR deleted by CRISPR/Cas9 (a gift from Dr. Jacqui Naylor, MedImmune) (49), were maintained in RPMI-1640 at 11 mM D-glucose, supplemented with 10% FBS, 10 mM HEPES, 2 mM L-glutamine, 1 mM pyruvate, 50 µM β-mercaptoethanol and 1% penicillin/streptomycin. MIN6B1 cells (a gift from Prof. Philippe Halban, University of Geneva, Switzerland) were maintained in DMEM at 25 mM D-glucose supplemented with 15% FBS, 50 µM β-mercaptoethanol and 1% penicillin/streptomycin. MIN6B1 cells with deleted endogenous GLP-1R expression were generated by CRISPR/Cas9 as follows: cells were co-transfected with the CRIPSR/Cas9 vector pX330-U6-Chimeric_BB-CBh-hSpCas9 (a gift from Prof. Feng Zhang, Addgene plasmid #42230) with cloned guide RNA sequence 5’-CCCCGAGCAGCAGGAGCGCC-3’ targeting the minus strand of mouse GLP-1R exon 1 and pcDNA 3.1+ and selected in 1 mg/ml G418. A mixed population of cells was recovered and cultured without further G418 selection. Stable SNAP-GLP-1R-expressing HEK293 cells [wild-type and β-arrestin-less (42)] were generated by transfection of pSNAP-GLP-1R (Cisbio) and G418 (1 mg/ml) selection. The same approach was used to generate HEK293 cells stably expressing N-terminal FLAG-tagged GLP-1R, and to reintroduce stable expression of wild-type or C438A mutant SNAP-GLP-1R into INS-1 832/3 cells lacking endogenous GLP-1R expression. Stable SNAP-GLP-1R CHO-K1 and MIN6B1 cells were described previously (19). Mycoplasma testing was performed yearly.

### Generation of SNAP-GLP-1R C438A and ProLink-tagged SNAP-GLP-1R wild-type and C438A vectors

SNAP-GLP-1R C438A mutant vector was generated from wild-type SNAP-tagged human GLP-1R (Cisbio) by site-directed mutagenesis with the QuikChange II XL Site-Directed Mutagenesis Kit (Agilent), following the manufacturer’s instructions. ProLink (PK)-tagged variants of both SNAP-GLP-1R wild-type and C438A vectors were generated for β-arrestin-2 recruitment assays by cloning of HindIII - BglII restriction fragments onto the pCMV-ProLink™ 1 Vector (DiscoverX).

### MβCD treatments

Cells were treated with MβCD (Sigma Aldrich) at the indicated concentration in HBSS, followed by washing. Where relevant, MβCD treatments were performed after labeling with SNAP-Surface fluorescent probes.

### Biochemical quantification of cellular cholesterol

Cells were treated with or without MβCD as above. After washing, lipid was extracted using butanol as described (72). Briefly, the washed cell pellet was left in 250 µl butanol overnight at 4°C, and the organic layer was aspirated, placed in a microplate and allowed to evaporate to dryness. The lipid film was re-dissolved in 1% Triton X-100 and cholesterol concentration determined using the Amplex Red Cholesterol Assay Kit (Thermo Fisher), followed by normalization to protein content of the cell pellet determined by BCA assay.

### Cell labeling for electron microscopy (EM)

SNAP-GLP-1R-expressing cells cultured on Thermanox coverslips (Agar Scientific) were labeled with 2 µM cleavable SNAP-Surface biotin probe in full media, followed by 5 µg/ml NaN_3_-free Alexa Fluor 488 Streptavidin, 10 nm colloidal gold conjugate (Thermo Fisher) in HEPES-bicarbonate buffer (120 mM NaCl, 4.8 mM KCl, 24 mM NaHCO_3_, 0.5 mM Na_2_HPO_4_, 5 mM HEPES, 2.5 mM CaCl_2_, and 1.2 mM MgCl_2_, saturated with 95% O_2_ / 5% CO_2_, pH 7.4) and 1% BSA, and stimulated with the indicated treatment. Conventional EM was performed as described. Briefly, cells were fixed, processed, mounted on Epon stubs, polymerized at 60 °C, and 70 nm sections cut *en face* with a diamond knife (DiATOME) in a Leica Ultracut UCT ultramicrotome before examination on an FEI Tecnai G2-Spirit TEM. Images were acquired in a charge-coupled device camera (Eagle), and processed in Fiji.

### EM quantification of clustering by plasma membrane rip-off

Cells expressing SNAP-GLP-1R were cultured on Thermanox coverslips, and gold-labeled with SNAP-Surface biotin followed by Alexa Fluor 488 Streptavidin, 10 nm colloidal gold conjugate as above. Cells were stimulated with the indicated treatments for 1 minute. Membrane rip-offs were performed following the protocol of Sanan and Anderson (73). In brief, EM grids were coated with formvar and poly-L-lysine and cell coverslips were inverted on top with pressure applied for ∼10 sec at 4°C. This allowed detachment of the apical cellular membranes onto the grids, which were immediately fixed by incubating in 4% glutaraldehyde in 25 mM HEPES buffer for 15 minutes. The grids were subsequently prepared for EM analysis by post-fixation in 2% aqueous osmium followed by 1% aqueous tannic acid and 1% aqueous uranyl acetate, for 10 minutes each at room temperature, and examined on a JEOL 1400+ TEM with images taken on an AMT digital camera. Gold particle nearest neighbor analysis was performed using ImageJ.

### Cell labeling for confocal and Total Internal Reflection Fluorescence (TIRF) microscopy

For confocal microscopy, cells were labeled at 37°C with 1 µM of the indicated SNAP-tag fluorescent probe (New England Biolabs) in full media, stimulated with agonists for the indicated times, fixed in 4% paraformaldehyde, mounted in Prolong Diamond antifade reagent with 4,6-diamidino-2-phenylindole (Life Technologies), and imaged with a Zeiss LSM-780 inverted confocal laser-scanning microscope in a 63x/1.4 numerical aperture oil-immersion objective from the Facility for Imaging by Light Microscopy (FILM) at Imperial College London, and analyzed in Fiji.

For TIRF microscopy, SNAP-GLP-1R-expressing cells were transfected with µ2-HA-WT (a gift from Prof. Alexander Sorkin, Addgene plasmid #32752), plated in glass bottom MatTek dishes, labeled with SNAP-Surface 488 probe as above, stimulated or not for 2 minutes with the indicated agonist, and fixed as above before immunofluorescence with a mouse monoclonal anti-HA (HA-7) antibody, catalogue no. H3663 (Sigma), followed by a secondary AlexaFluor 546 antibody. Un-mounted samples in PBS were imaged using a Nikon Eclipse Ti microscope equipped with a ×100/1.49 numerical aperture TIRF objective, a TIRF iLas2 module and a Quad Band TIRF filter cube (TRF89902, Chroma). Images were acquired with an ORCA-Flash 4.0 camera (Hamamatsu) and Metamorph software (Molecular Devices), and analyzed in Fiji.

### Clustering measurements by TIRF-Photoactivated Localization Microscopy (PALM)

For TIRF-PALM analysis of FLAG-GLP-1R cluster sizes, the GLP1R-Tango vector (a gift from Prof Bryan Roth, Addgene plasmid #66295), which harbors a single N-terminal FLAG tag, was used to introduce a STOP codon at the end of the GLP-1R coding sequence by site-directed mutagenesis performed as above. HEK293 cells were used to generate a subline with stable expression of the engineered FLAG-GLP-1R construct as above. Cells were subsequently plated onto glass bottom MatTek dishes and labeled with the anti-FLAG GAGE 500 antibody (1:500 dilution) described in (28) in 10% FCS in PBS containing calcium and magnesium at 37°C for 30 minutes. Cells were washed in PBS containing calcium and magnesium and stimulated or not with 100 nM exendin-4 for 2 minutes prior to fixation for 30 minutes in 4% paraformaldehyde with 0.2% glutaraldehyde to minimize antibody-induced clustering artifacts. Following fixation, cells were washed in PBS with calcium and magnesium and maintained in the same buffer for imaging. All steps were carried out in the dark to ensure minimal label photo-switching. Images were acquired using a Zeiss Elyra PS1 featuring an AxioObserver Z1 motorized inverted microscope with TIRF capability with an Alpha Plan-APO 1.46 numerical aperture ×100 oil immersion objective at the Imperial College London FILM facility. Photo-conversion of CAGE 500 was achieved with a 405 nm light source and simultaneously imaged and photo-bleached with a 488 nm laser line. The microscope was contained in a plastic draft-proof enclosure maintained at a constant temperature of 25°C and mounted on a vibration isolation table. Laser lines were switched on at least 1 hour prior to imaging to allow stabilization of the system in order to ensure minimal sample drift throughout the imaging. Each TIRF-PALM time-series was acquired using a cooled electron multiplying charged coupled device camera (EM-CCD; iXon DU 897, Andor Technology) and LMS Zen operating software with an exposure time of 30 ms. Localization of receptors was determined using QuickPALM Fiji plugin: fluorescent images of cropped non-overlapping areas of 7 × 7 µm within cell borders were analyzed using a full-width half-maximum value of 4, and signal to noise ratio of 8. Analyzed areas did not span cell membranes to exclude any potential biasing resulting from edge effects. Data tables containing particle localization 2-D coordinates were generated. The number of associated receptor molecules from these coordinates was determined with PD-Interpreter as in (28) with a 50 nm radius from each fluorophore. To discount any overestimation of oligomers, events within a 10 nm radius of an activated fluorophore were discounted from the analysis.

### Clustering measurements by Homogeneous Time Resolved Fluorescence (HTRF)

Cells stably or transiently expressing SNAP-GLP-1R were dual-labeled with the TR-FRET probe SNAP-Lumi4-Tb (40 nM, Cisbio) and SNAP-Surface 647 (1 µM, New England Biolabs) for 60 minutes at room temperature. These concentrations were selected as they provided optimum signal intensity at both measurement wavelengths. Where relevant, MβCD treatments were performed after labeling. Cells were washed and placed in HBSS in a white plate for a 10 minutes baseline measurement at 37°C using a Spectramax i3x (Molecular Devices) plate reader fitted with a TR-FRET filter set (λEx = 335 nm, λEm = 616 nm and 665 nm, delay 30 µs, integration time 400 µs). TR-FRET was sequentially measured after agonist addition. The ratio of fluorescent signals at both emission wavelengths (665 / 616) was considered indicative of clustering as it reflects both transient and stable interactions between receptor protomers occurring within the long fluorescence lifetime of the excited terbium cryptate. For agonist comparisons within the same cell type, individual well responses were normalized to baseline to reduce variability; where different cell types, receptor mutants, or pre-treatments were compared, results were expressed as the non-normalized ratio so that baseline differences could be observed. Kinetic data were found to fit a 4-parameter logistic curve, and this approach was used to quantify half times.

### NR12S assays and analysis

Cells expressing SNAP-GLP-1R were first labeled with SNAP-Lumi4-Tb (40 nM) for 60 minutes at room temperature. Where relevant, MβCD treatments were performed after Lumi4-Tb labeling. Spectral measurements and analysis: 1) Washed, Lumi4-Tb-labeled cells were placed in HBSS in a white plate and a full baseline TR-FRET spectral scan was performed at 37°C using a Flexstation 3 plate reader (Molecular Devices; λEx = 335 nm, λEm = 460 – 650 nm in 5 nm steps, no cutoff, delay 50 µs, integration time 300 µs) to detect the Lumi4-Tb-only signal; 2) freshly-prepared NR12S in HBSS was then added to the desired concentration, and after a further 5 minutes, a second spectral scan was performed to detect basal FRET; 3) agonist was added for 5 minutes and a final spectral scan was performed to detect agonist-changes in TR-FRET. After an initial blank subtraction, data were analyzed by ratiometrically normalizing the spectrum from each read to the signal measured at 490 nm; the normalized Lumi4-Tb-only read was subtracted from subsequent normalized reads to obtain the NR12S-specific TR-FRET signal in the presence and absence of agonist. Where indicated, the area under the curve (AUC) was calculated below and above the λEm maximum of 590 nm as a marker of relative contributions from liquid-ordered (530 - 590 nm) and liquid-disordered (590 - 650 nm) -associated SNAP-GLP-1Rs, and expressed relative to the total NR12S TR-FRET AUC (530 - 650 nm). Alternatively, liquid-ordered and liquid-disordered FRET were assigned values of 570 and 610 nm respectively, with the NR12S-associated increase at each wavelength ratiometrically-expressed to indicate differences in SNAP-GLP-1R localization in each nanodomain. Kinetic measurements and analysis: Washed, Lumi4-Tb-labeled cells were treated with NR12S at the indicated concentration and a 5-minute baseline read was performed (λEx = 335 nm, λEm = 490, 570 and 610 nm). Agonist was then added and changes in TR-FRET sequentially monitored. Signals at each wavelength were blank-subtracted and expressed ratiometrically to each other, with agonist-induced changes quantified relative to individual well baseline.

### Binding kinetics measurements

Cells expressing SNAP-GLP-1R were first labeled with SNAP-Lumi4-Tb (40 nM) for 60 minutes at room temperature. Where relevant, MβCD treatments were performed after Lumi4-Tb labeling. After washing, cells were placed in HBSS + 0.1% BSA, supplemented with metabolic inhibitors (20 mM 2-deoxyglucose and 10 mM NaN_3_) to prevent GLP-1R endocytosis (74), and baseline TR-FRET was measured at over 10 minutes at 37°C using a Flexstation 3 plate reader (λEx = 335 nm, λEm = 520 and 620 nm, delay 50 µs, integration time 300 µs). FITC-conjugated agonists were then added and TR-FRET sequentially measured to monitor agonist association. Binding was quantified ratiometrically as 520 / 620 signal, and data were analyzed using the GraphPad Prism model “association kinetics: two or more concentrations of hot” to obtain association (*k*_on_) and dissociation (*k*_off_) rate constants, from which the equilibrium binding constant K_d_ was determined.

### HTRF cAMP measurements

Cells were treated in white microplates with agonist, with or without IBMX to inhibit phosphodiesterases, for the indicated period of time, before total cellular cAMP determination by HTRF (Cisbio cAMP Dynamic 2).

### Measurement of β-arrestin-2 recruitment by PathHunter assay

Cells were treated with agonist or drug at the indicated concentration for the indicated time periods before lysis and/or membrane fractionation and detection of β-arrestin-2 recruitment by the PathHunter enzyme fragment complementation assay (DiscoverX) according to the manufacturer’s instructions.

### Real-time measurement of cAMP production and PKA activation using FRET biosensors

The ^T^Epac^VV^ biosensor was a gift from Dr K Jalink, The Netherlands Cancer Institute. The AKAR4-NES and AKAR4-Lyn biosensors were gifts from Dr Jin Zhang (Addgene plasmid #61620). Cells expressing SNAP-GLP-1Rs were transiently transfected with the relevant biosensor plasmid for 36 hours prior to the assay and placed in HBSS in 96-well clear bottom black plates. After a 5 minute baseline read at 37°C (λEx = 440 nm, λEm = 485 and 535 nm) in a Flexstation 3 plate reader, compounds were injected to each well and sequential reads immediately commenced. FRET was expressed ratiometrically as signal at 535 nm divided by signal at 485 nm. For agonist comparisons within the same cell type, individual well responses were normalized to baseline to reduce variability; where different cell types, receptor mutants or pre-treatments were compared, results were expressed as the non-normalized ratio so that baseline differences could be observed. Dose-responses were either analyzed by obtaining the cumulative AUC relative to baseline over a fixed time period, or at each time point to allow kinetic comparisons of agonists between each pathway.

### Measurement of GLP-1R internalization by DERET assay

The assay was performed as previously described (19). Cells expressing SNAP-GLP-1R were first labeled with SNAP-Lumi4-Tb (40 nM) for 60 minutes at room temperature. Where relevant, MβCD treatments were performed after Lumi4-Tb labeling. After washing, cells were placed in HBSS supplemented with 24 mM fluorescein. A 10 minute baseline read at 37°C was taken in a Flexstation 3 plate reader in HTRF mode (λEx = 335 nm, λEm = 520 and 620 nm, delay 400 µs, integration time 1500 µs), after which compounds were injected to the wells. Loss of surface SNAP-GLP-1R was detected as changes over time in the HTRF ratio (signal at 620 nm divided by that at 520 nm, after blank subtraction at each time point). Half times were obtained via 4-parameter logistic curve fitting.

### Insulin secretion assays

INS-1 832/3 cells with endogenous GLP-1R expression deleted by CRISPR/Cas9 and stably expressing wild-type or C438A SNAP-GLP-1R were pre-incubated overnight in 3 mM glucose medium. Exendin-4 was added in complete medium at 11 mM glucose at the time of seeding into plates. After 16 h incubation, samples were obtained for secreted insulin, and analyzed by HTRF (Cisbio). Results were normalized as a fold increase compared to cells treated with 11 mM glucose alone.

### Palmitoylation assays

For the detection of palmitoylated GLP-1Rs, cells were stimulated with 100 nM of the indicated agonist for 10 minutes and, when appropriate, pre-treated with 10 µM BETP for 5 minutes (and throughout the agonist stimulation period) or overnight with 200 µM of the palmitoylation inhibitor 2-bromopalmitate. Isolation of palmitoylated proteins was performed with the CAPTUREome™ S-Palmitoylated Protein Kit (Badrilla), following the manufacturer’s instructions. The assay is based on the acyl-resin assisted capture method, described in detail elsewhere (75). Key steps include blocking of free thiol groups with a blocking reagent, cleavage of thioester linkages to release the palmitate group, and capture of newly liberated thiols with the CAPTUREome™ capture resin. Briefly, cells were lysed and free thiols blocked in the manufacturer’s blocking buffer at 40°C for 4 hours with constant vortexing. Proteins were then precipitated with cold acetone for 20 minutes at -20°C. Following centrifugation at 16,000 g for 5 minutes, the pellet was extensively washed with 70% cold acetone and resuspended in the manufacturer’s binding buffer. Approximately 10% of the soluble lysate was saved as the input fraction (IF). Prewashed CAPTUREome™ capture resin was added to the lysates. To this mixture, either the thioester cleavage or the acyl-preservation reagents were added. Resins were washed five times with the manufacturer’s binding buffer. Urea sample buffer (100 mM Tris-HCl pH 6.8, 2.5% SDS, 4 M urea, 50 mM dithiothreitol, 0.05% bromophenol blue) was used to elute captured proteins from the resin. Eluted samples were incubated at 37°C for 30 minutes before Western blotting for SNAP-GLP-1R to detect the fraction of palmitoylated *versus* non-palmitoylated receptor. Resin-bound proteins treated with the thioester cleavage reagent are referred to as the cleaved bound fraction (cBF), and represent the fraction of palmitoylated proteins, while resin-bound proteins treated with the acyl-preservation reagent are referred to as the preserved bound fraction (pBF) and are used as negative control for the assay. Corresponding unbound fractions are known as the cleaved unbound fraction (cUF) and the preserved unbound fraction (pUF).

### Detergent-resistant and detergent-soluble membrane fractionation

For membrane raft purification experiments, the cells were seeded onto 10-cm dishes and transiently transfected with GαS-YFP (a gift from Catherine Berlot, Addgene plasmid #55781) when indicated. After the indicated treatments, cells were osmotically lysed in 20 mM Tris-HCl (pH 7.0) supplemented with cOmplete EDTA-free protease inhibitor cocktail (Roche). The cell suspension was homogenized by passing through a 22-gauge needle and ultracentrifuged at 63,000 g for 1 h at 4°C with a Sorvall Discovery M120 ultracentrifuge. The supernatant was discarded and the pellet was resuspended in ice-cold PBS supplemented with protease inhibitor cocktail and an aliquot retained for total membrane fraction (TMF) analysis. PBS was added to a final volume of 1 ml and the suspensions were ultracentrifuged at 63,000 g for 30 minutes at 4°C. The supernatant was discarded, and the pellet resuspended in ice-cold PBS supplemented with 1% Triton-X100 and protease inhibitor cocktail, and incubated for 30 minutes at 4°C. Samples were ultracentrifuged again at 63,000 g for 1 h at 4°C. The supernatant, containing the detergent-soluble membrane fraction (DSF), was retained for analysis, and the detergent-resistant membrane fraction (DRF) pellet was resuspended in 1% SDS with protease inhibitor cocktail. All the samples were sonicated with the Ultrasonic cell disruptor XL-2000 probe sonicator (Misonix Inc.) and centrifuged at 14,000 g for 5 minutes at 4°C before analysis by SDS-PAGE and Western blotting. For measurement of β-arrestin-2 recruitment to different membrane regions, purified fractions from PathHunter CHO-GLP-1R cells prepared as above after vehicle / exendin-4 treatment were diluted in HBSS and measured using PathHunter chemiluminescent reagents.

### SDS-PAGE and Western blotting

Samples were separated by SDS-PAGE on 10% gels under reducing conditions, immunoblotted onto PVDF membranes (GE Healthcare) and bands detected by enhanced chemiluminescence (GE Healthcare) onto films developed on a Xograph Compact X5 processor. Antibodies for Western blotting were: anti-flotillin-1 mouse monoclonal antibody, catalogue no. sc-74566, Santa Cruz Biotechnology (1:200 dilution); anti-SNAP-tag rabbit polyclonal antibody, catalogue no. P9310S, New England Biolabs (1:500 dilution); anti-GFP mouse monoclonal antibody, catalogue no. G6795, Sigma (1:1000 dilution); anti-human transferrin receptor mouse monoclonal antibody, catalogue no. G1/221/12, Developmental Studies Hybridoma Bank (DSHB; 0.2 µg/ml); anti-human GLP-1R mouse monoclonal antibody, catalogue no. Mab 3F52, DSHB (0.2 µg/ml); anti-mouse GLP-1R mouse monoclonal antibody, catalogue no. Mab 7F38, DSHB (0.2 µg/ml); and anti-α-tubulin mouse monoclonal antibody, catalogue no. T5168, Sigma (1:1000 dilution).

### Statistical analysis

Statistical significance was assessed using Student’s *t* test or ANOVA as indicated, with matched analyses performed where possible. Statistical analyses were performed using GraphPad Prism 7.0. Statistical significance was taken as p<0.05. Unless indicated, values represented are the mean ± SEM.

## Acknowledgements

This work was funded by MRC project grant MR/R010676/1. MIN6B1 cells were kindly provided by Prof. Philippe Halban (University of Geneva, Switzerland) with permission from Prof. Jun-ichi Miyazaki (University of Osaka, Japan) who produced the maternal MIN6 cell line. INS-1 832/3 cells were kindly provided by Prof. Christopher Newgard (Duke University, USA), and INS-1 832/3 cells with endogenous GLP-1R or GIPR deleted by CRISPR/Cas9 were kindly provided by Dr Jacqueline Naylor at MedImmune (AstraZeneca).

## Supplementary Figure Legends

**Figure S1.**
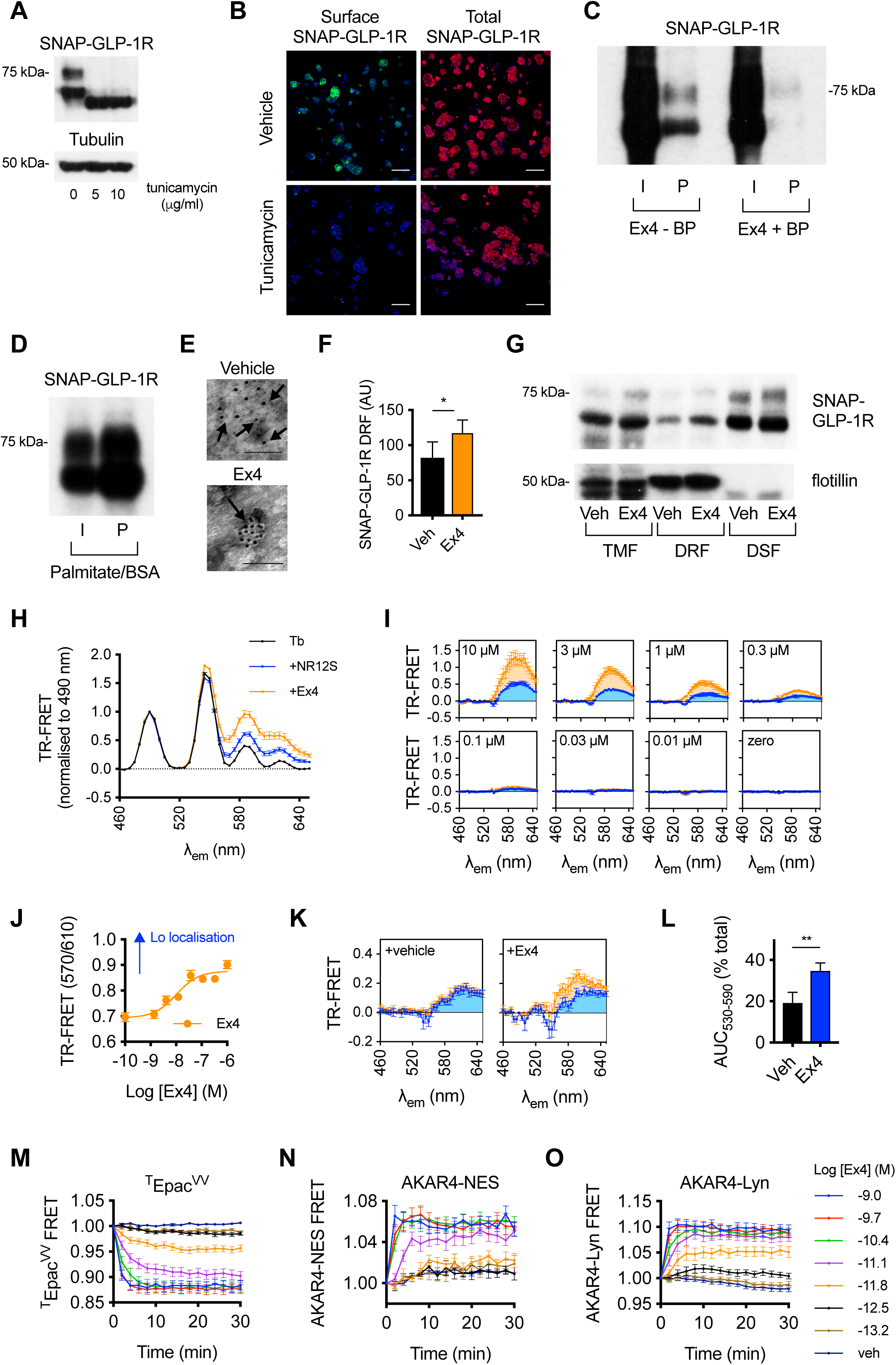
Agonist-induced GLP-1R palmitoylation and clustering in membrane nanodomains – extra data. (**A**) Western blotting analysis of SNAP-GLP-1R in lysates from CHO-K1 cells stably expressing SNAP-GLP-1R incubated overnight with the indicated amount of tunicamycin, a drug that blocks N-linked glycosylation, showing that the two predominant bands detected for SNAP-GLP-1R at ∼75 and ∼65 kDa correspond to differentially glycosylated forms of the receptor. Note that an additional lower band corresponding to non-glycosylated receptor is occasionally detected for total SNAP-GLP-1R (Figure 1A). Tubulin is shown as loading control. (**B**) Confocal analysis of surface and total SNAP-GLP-1R levels in MIN6B1 cells stably expressing SNAP-GLP-1R following overnight incubation in vehicle or 10 µg/ml tunicamycin. Surface SNAP-GLP-1R labeled with SNAP-Surface 488, green; total SNAP-GLP-1R labeled with SNAP-Cell TMR-Star, red; nuclei (DAPI), blue; size bars, 100 µm. (**C**) Total input (I) and palmitoylated (P) SNAP-GLP-1R fractions from MIN6B1 cells stably expressing SNAP-GLP-1R and treated with 100 nM exendin-4 (Ex4) for 10 min, with and without pre-treatment with 200 µM bromopalmitate (BP) overnight. (**D**) Total input (I) and palmitoylated (P) SNAP-GLP-1R fractions from MIN6B1 cells stably expressing SNAP-GLP-1R and treated with 200 µM palmitate/BSA overnight. (**E**) Representative images of gold-labeled SNAP-GLP-1Rs (arrows) from EM analysis of 2-D plasma membrane sheets isolated from MIN6B1 beta cells stably expressing SNAP-GLP-1R following gold labeling of SNAP-tagged receptors and treatment with vehicle (Veh) or 100 nM exendin-4 for 2 min. (**F**) Quantification of SNAP-GLP-1R levels in DRF fractions of vehicle-and exendin-4-treated MIN6B1 cells stably expressing SNAP-GLP-1R, *n*=3, paired *t*-test. (**G**) SNAP-GLP-1R distribution within total membrane (TMF), detergent-resistant (DRF) and detergent-soluble (DSF) fractions isolated from HEK293 cells stably expressing SNAP-GLP-1R treated with vehicle or 100 nM exendin-4 for 2 min, with flotillin as a marker of membrane raft enrichment. (**H**) TR-FRET spectra in Lumi4-Tb-labeled HEK-SNAP-GLP-1R cells treated sequentially with 1 µM NR12S and exendin-4 (100 nM), normalized to signal at 490 nm, *n*=3. (**I**) As for (H) but after subtraction of Lumi4-Tb-only signal, with each panel indicating a different concentration (indicated) of NR12S applied. (**J**) Alternative analysis of data shown in Figure 1G-I, with TR-FRET increase after Lumi4-Tb-only subtraction quantified at 570 nm and 610 nm and expressed ratiometrically to indicate increased localization of SNAP-GLP-1R in Lo phase, 3-parameter fit shown. (**K**) TR-FRET spectra in Lumi4-Tb-labeled monoclonal CHO-SNAP-GLP-1R cells treated sequentially with NR12S (1 µM) and either vehicle or exendin-4 (100 nM), normalized to signal at 490 nm, *n*=4. (**L**) Quantification of Lo-associated SNAP-GLP-1R-NR12S TR-FRET with and without exendin-4 treatment, determined from (K) as percentage of total AUC from 530 - 590 nm portion of spectrum, paired *t*-test. (**M**) FRET traces at the indicated concentration of exendin-4 in CHO-SNAP-GLP-1R cells expressing ^T^Epac^VV^ to measure cellular cAMP, expressed ratiometrically as signal at 535 nm divided by signal at 485 nm and then normalized to individual well baseline, *n*=15. (**N**) As for (M) but with AKAR4-NES biosensor to measure cytosolic PKA activation, *n*=5. (**O**) As for (M) but with AKAR4-Lyns biosensor to measure raft-associated PKA activation, *n*=11. Data are depicted as mean ± SEM; **p<0.01, by statistical test indicated in the text.

**Figure S2.**
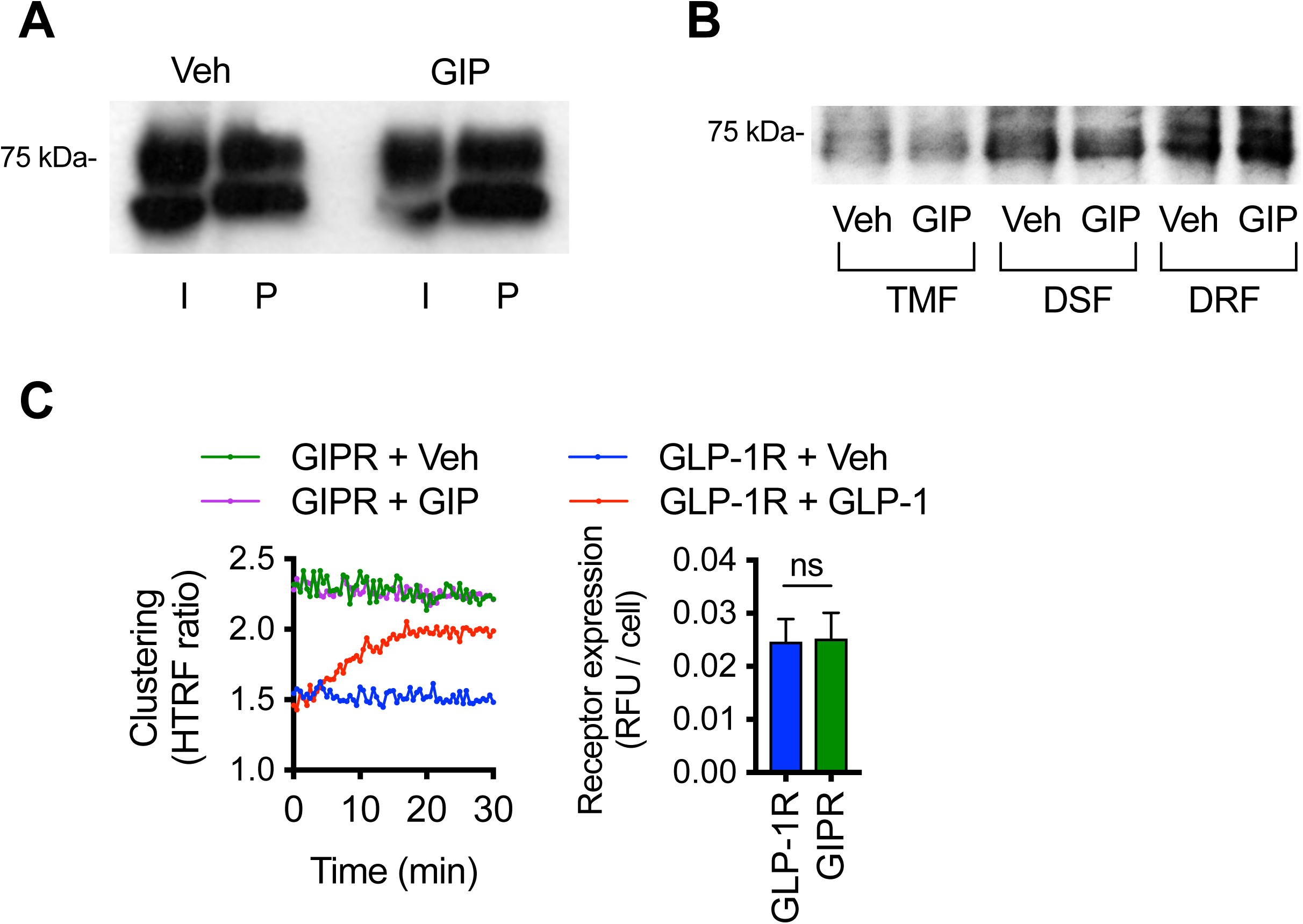
GIPR palmitoylation and membrane nanodomain clustering responses. (**A**) Total input (I) and palmitoylated (P) SNAP-GIPR fractions from CHO-K1 cells stably expressing SNAP-GIPR and treated with vehicle (Veh) or 100 nM GIP for 10 min. (**B**) SNAP-GIPR distribution within total membrane (TMF), detergent-soluble (DSF) and detergent-resistant (DRF) fractions isolated from INS1 832/3 cells with endogenous GIPR expression deleted by CRISPR/Cas9 and expressing SNAP-GIPR after treatment with vehicle or 100 nM GIP for 5 min. (**C**) Comparison of SNAP-GLP-1R and SNAP-GIPR clustering, detected by HTRF, in transiently-transfected HEK293 cells treated with vehicle, GLP-1 (100 nM), or GIP (100 nM) as indicated, *n*=3, error bars have been omitted for clarity. Parallel measurement of receptor surface expression by Lumi4-Tb labeling and normalization to cell count also shown, *n*=3, non-significant (ns), paired *t*-test. Unless stipulated, data are depicted as mean ± SEM; ns=non-significant, by statistical test indicated in the text.

**Figure S3.**
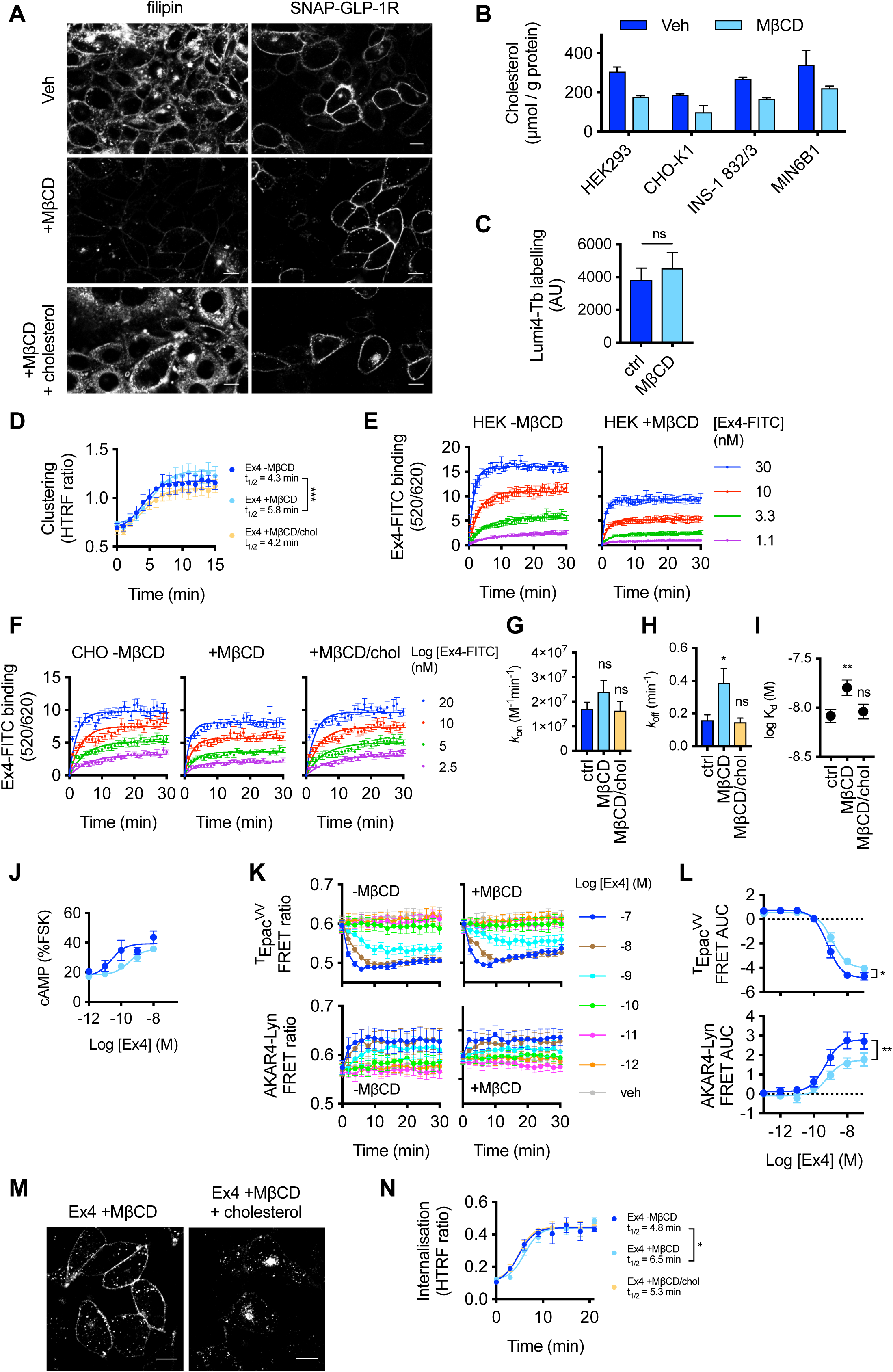
Effects of cholesterol depletion by MβCD treatment on SNAP-GLP-1R clustering, membrane nanodomain recruitment, signaling and internalization – extra data. (**A**) Cholesterol levels determined by filipin staining (left) in CHO-K1 cells stably expressing SNAP-GLP-1R after SNAP-Surface 549 labeling (right) and treated with vehicle (Veh), MβCD (10 mM) or MβCD saturated with cholesterol for 1 hour as a control. Filipin staining was carried out with the Cell-Based Cholesterol Assay Kit (Abcam). Size bars, 10 µm. (**B**) Biochemical quantification of cholesterol depletion by MβCD in HEK293, CHO-K1, INS-1 832/3 and MIN6B1 cells treated for 45 min with 10 mM MβCD (3 mM for CHO-K1 cells) followed by butanol extraction and cholesterol quantification and normalization to protein content, *n*=3. (**C**) Lack of effect of MβCD (10 mM, 45 min) treatment on surface labeling by Lumi4-Tb in HEK-SNAP-GLP-1R cells; measured as TR-FRET at 550 nm and normalized for cell count, *n*=3. (**D**) Exendin-4 (Ex4)-induced (100 nM) SNAP-GLP-1R clustering measured by HTRF in monoclonal CHO-SNAP-GLP-1R cells with and without prior treatment with MβCD (3 mM, 45 min), MβCD saturated with cholesterol, or vehicle, expressed as HTRF ratio, *n*=5, 4-parameter logistic fit shown and used to quantify t_1/2_, compared by paired *t*-test. (**E**) Kinetic binding response of exendin-4-FITC (Ex4-FITC) measured by TR-FRET in Lumi4-Tb-labeled HEK-SNAP-GLP-1R cells, with or without treatment with MβCD (10 mM, 45 min), expressed ratiometrically as 520 / 620 nm signal after blank subtraction, *n*=4. Relates to Figure 2D-F. (**F**) Kinetic binding of exendin-4-FITC (Ex4-FITC) measured by TR-FRET in Lumi4-Tb-labeled CHO-SNAP-GLP-1R cells, following treatment with MβCD (3 mM, 45 min), MβCD saturated with cholesterol, or vehicle, expressed ratiometrically as 520 / 620 nm signal after blank subtraction, *n*=9. (**G**) Association rate constant (*k*_on_) determined from (F), one-way randomized block ANOVA with Dunnett’s test *vs.* control treatment. (**H**) Dissociation rate constant (*k*_off_) determined from (F), one-way randomized block ANOVA with Dunnett’s test *vs.* control treatment. (**I**) Equilibrium binding constant (K_d_) determined from (F), one-way randomized block ANOVA with Dunnett’s test *vs.* control treatment. (**J**) Effect of MβCD (10 mM, 45 min) on exendin-4-induced cAMP production in MIN6B1 cells, 10 min stimulation with 500 µM IBMX, normalized to forskolin (FSK, 10 µM), *n*=4. (**K**) Effect of MβCD (10 mM, 45 min) treatment on exendin-4-induced ^T^Epac^VV^ and AKAR4-Lyn responses in stable HEK-SNAP-GLP-1R cells, expressed as HTRF ratio, *n*=5. (**L**) Dose responses for exendin-4-induced ^T^Epac^VV^ and AKAR4-Lyn FRET changes, determined as AUC over 30 min relative to individual baselines from data shown in (K), paired *t*-test used to compare E_max_. (**M**) Confocal analysis of SNAP-GLP-1R internalization in CHO-K1 cells stably expressing SNAP-GLP-1R and labeled with SNAP-Surface 549 prior to treatment with MβCD (10 mM) or MβCD saturated with cholesterol for 1 hour followed by 15 min stimulation with 100 nM exendin-4. (**N**) As for (M) but indicating SNAP-GLP-1R internalization as measured by DERET with and without MβCD pre-treatment (3 mM, 45 min), *n*=5, compared by paired *t*-test. Data are depicted as mean ± SEM; ns=non-significant, *p<0.05, **p<0.01, ***p<0.001, by statistical test indicated in the text.

**Figure S4.**
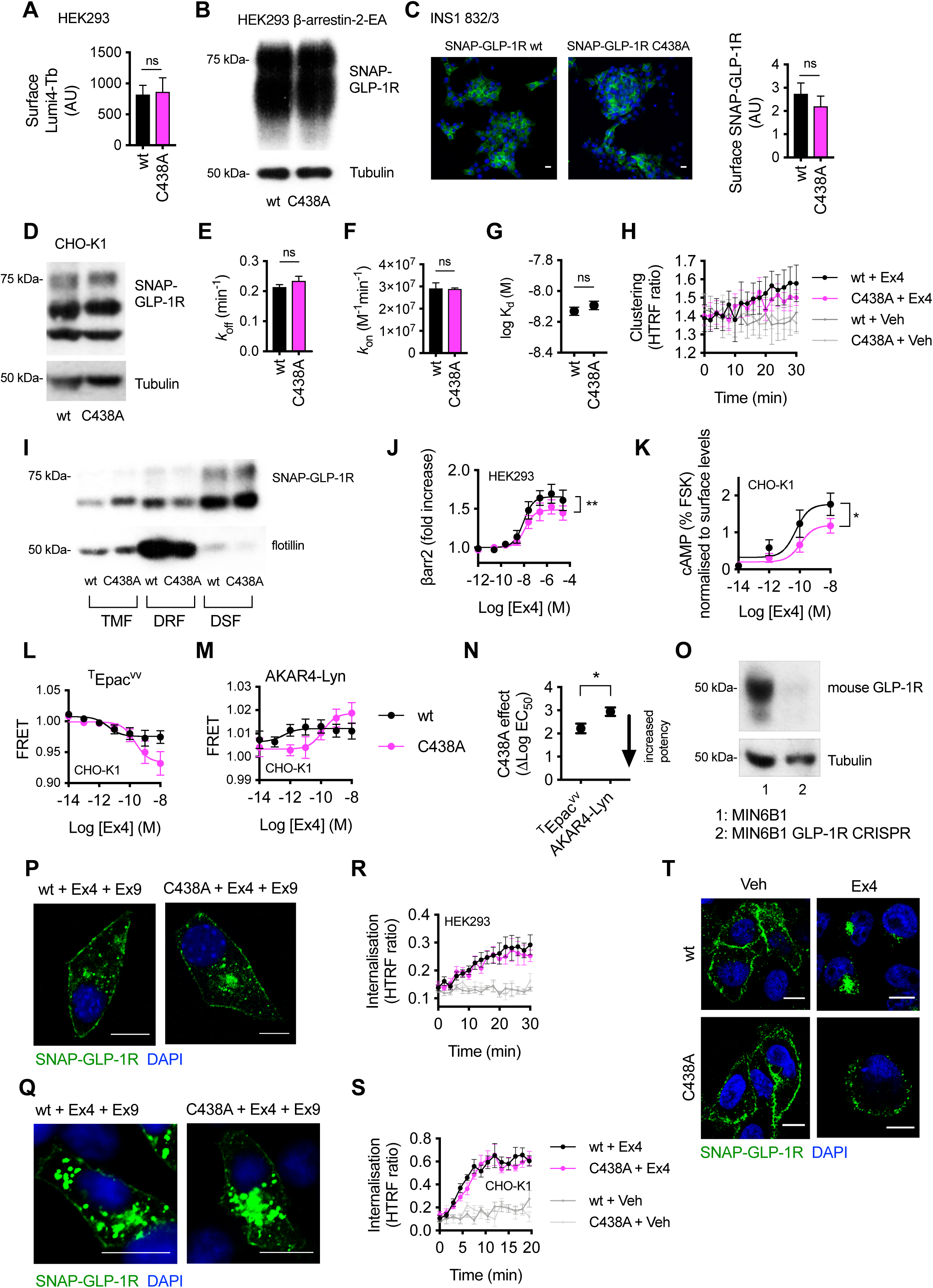
Effects of C438A on SNAP-GLP-1R palmitoylation, clustering, membrane nanodomain recruitment and signaling, and internalization – extra data. (**A**) Surface Lumi4-Tb labeling in HEK293 cells transiently transfected with wild-type (wt) or C438A SNAP-GLP-1R, *n*=9, paired *t*-test. (**B**) SNAP-GLP-1R wild-type or C438A expression levels in lysates from HEK293 β-arrestin-2-EA cells transiently transfected with wild-type or C438A SNAP-GLP-1R-PK constructs. Tubulin is shown as loading control. (**C**) SNAP-GLP-1R wild-type or C438A surface levels (right) and representative images (left) from INS1 832/3 cells with endogenous GLP-1R expression deleted by CRISPR/Cas9 and stably expressing either wild-type or C438A SNAP-GLP-1R. SNAP-Surface 488, green; nuclei (DAPI), blue; size bars, 10 µm; *n*=3, paired *t*-test. (**D**) SNAP-GLP-1R wild-type or C438A expression levels in lysates from CHO-K1 cells stably expressing SNAP-GLP-1R wild-type or C438A. Tubulin is shown as loading control. (**E-G**) Association rate constant (*k*_on_, E), dissociation rate constant (*k*_off_, F), and equilibrium binding constant (K_d_, G) expressed as log, derived from kinetic binding of exendin-4-FITC measured by TR-FRET in Lumi4-Tb-labeled CHO-K1 cells expressing SNAP-GLP-1R wild-type *vs.* C438A, paired *t*-test. (**H**) Exendin-4 (Ex4, 100 nM) -induced clustering of wild-type or C438A SNAP-GLP-1R transiently transfected in HEK293 cells, measured by HTRF, *n*=4. Relates to Figure 3D. (**I**) SNAP-GLP-1R wt *vs.* C438A distribution within total membrane (TMF), detergent-resistant (DRF) and detergent-soluble (DSF) fractions isolated from CHO-K1 cells stably expressing wild-type or C438A SNAP-GLP-1R and treated with 100 nM exendin-4 for 2 min. (**J**) Dose-response curves of β-arrestin-2 recruitment to the GLP-1R in HEK293 β-arrestin-2-EA cells transiently transfected with wild-type or C438A SNAP-GLP-1R-PK, normalized to basal response, *n*=5, paired *t*-test used to compare E_max_. (**K**) Exendin-4-induced cAMP dose-responses in CHO-K1 cells stably expressing wild-type or C438A SNAP-GLP-1R, normalized to forskolin (FSK, 10 µM) and receptor surface levels, *n*=4, 3-parameter fits shown, E_max_ compared by paired *t*-test. (**L**) Exendin-4-induced FRET responses in CHO-K1 cells stably expressing wild-type or C438A SNAP-GLP-1R and transiently transfected with ^T^Epac^VV^, normalized to individual well baseline, *n*=5, 3-parameter fit shown. (**M**) As for (L), but with AKAR4-Lyn. (**N**) Relative impact of the C438A mutation on total cAMP or lipid raft PKA signaling, determined (respectively) from Log EC_50_ values from ^T^Epac^VV^ or AKAR4-Lyn responses shown in (L) and (M) by subtracting C438A from wild-type result for each biosensor, paired *t*-test. (**O**) Endogenous GLP-1R level of expression in lysates from MIN6B1 wild-type *vs.* mouse GLP-1R CRISPR/Cas9-engineered cells. Tubulin is shown as a loading control. (**P**, **Q**) Confocal analysis of SNAP-GLP-1R wild-type *vs.* C438A plasma membrane recycling in MIN6B1 (P) and INS1 832/3 (Q) cells with endogenous GLP-1R expression deleted by CRISPR/Cas9 and expressing either wild-type or C438A mutant SNAP-GLP-1R following stimulation with 100 nM exendin-4 for 1 hour, wash-out and a further 3 hour incubation with 10 µM GLP-1R antagonist exendin(9-39). Nuclei (DAPI), blue; size bars, 10 µm. (**R**) Exendin-4 (100 nM) -induced internalization of wild-type or C438A SNAP-GLP-1R transiently transfected in HEK293 cells, measured by DERET, *n*=7. Relates to Figure 3J. (**S**) Exendin-4 (100 nM) -induced internalization of wild-type or C438A SNAP-GLP-1R stably transfected in CHO-K1 cells, measured by DERET, *n*=5. (**T**) Confocal analysis of SNAP-GLP-1R wild-type *vs.* C438A localization in stably expressing CHO-K1 cells following treatment with vehicle or 100 nM exendin-4 for 15 min. SNAP-Surface 488, green; nuclei (DAPI), blue; size bars, 10 µm. Data are depicted as mean ± SEM; ns=non-significant, *p<0.05, **p<0.01, by statistical test indicated in the text.

**Figure S5.**
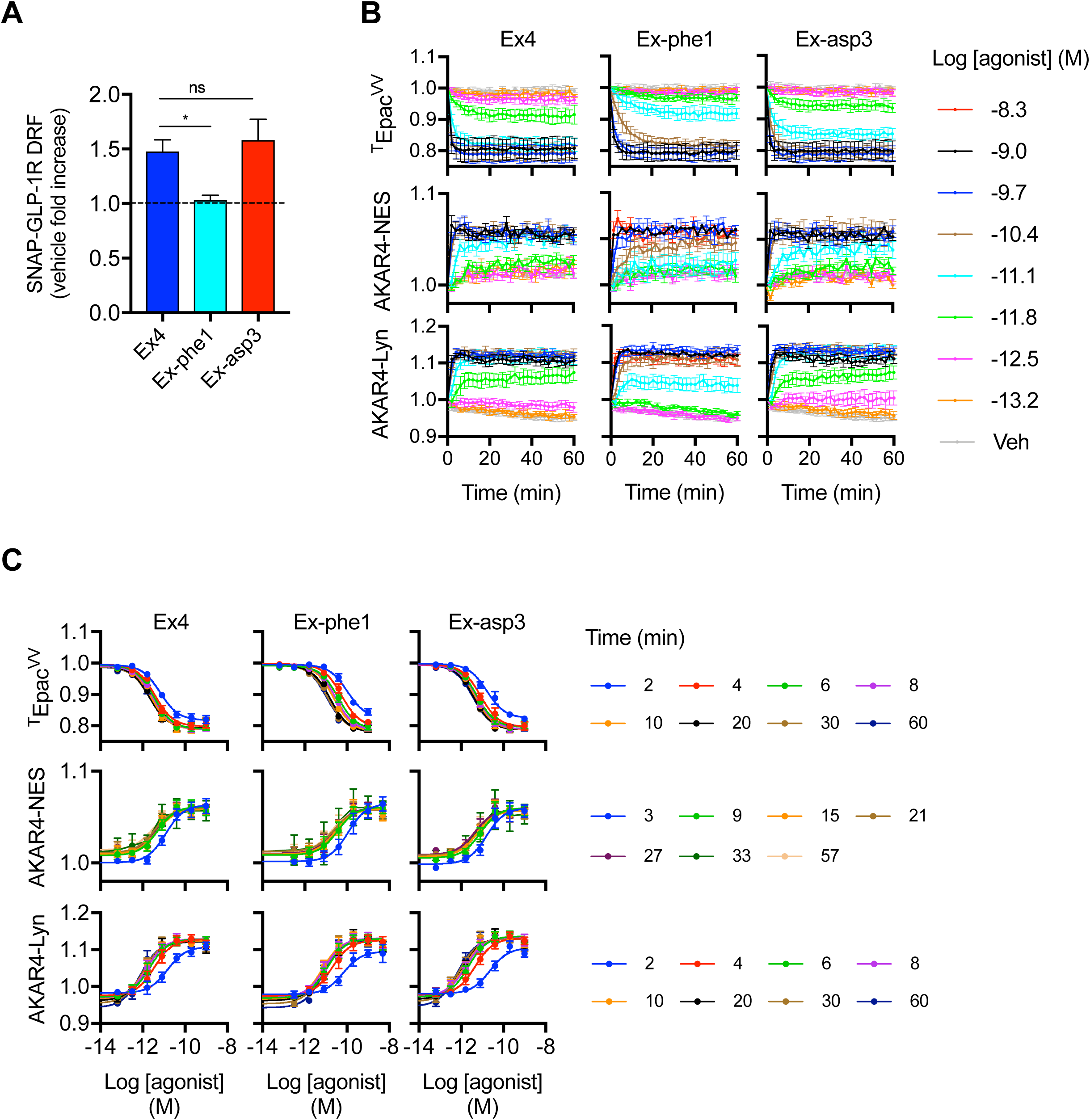
Biased agonist effect on SNAP-GLP-1R recruitment to and signaling from membrane nanodomains – extra data. (**A**) Quantification of SNAP-GLP-1R levels in DRF fractions of the indicated agonist in MIN6B1 cells stably expressing SNAP-GLP-1R, normalized to vehicle, *n*=4, one-way ANOVA with Dunnett’s test *vs.* exendin-4. (**B**) FRET traces with the indicated biosensors and agonists, normalized to individual well baseline, in CHO-SNAP-GLP-1R cells, *n*=5 for each. (**C**) Dose-response curves constructed from data shown in (B) for ^T^Epac^VV^, AKAR4-NES, and AKAR4-Lyn, for AKAR4-NES signals have been combined into 6-min bins to improve precision and named according to the mid-point of each period, 3-parameter fit shown. Data are depicted as mean ± SEM; ns=non-significant, *p<0.05, **p<0.01, by statistical test indicated in the text.

**Figure S6.**
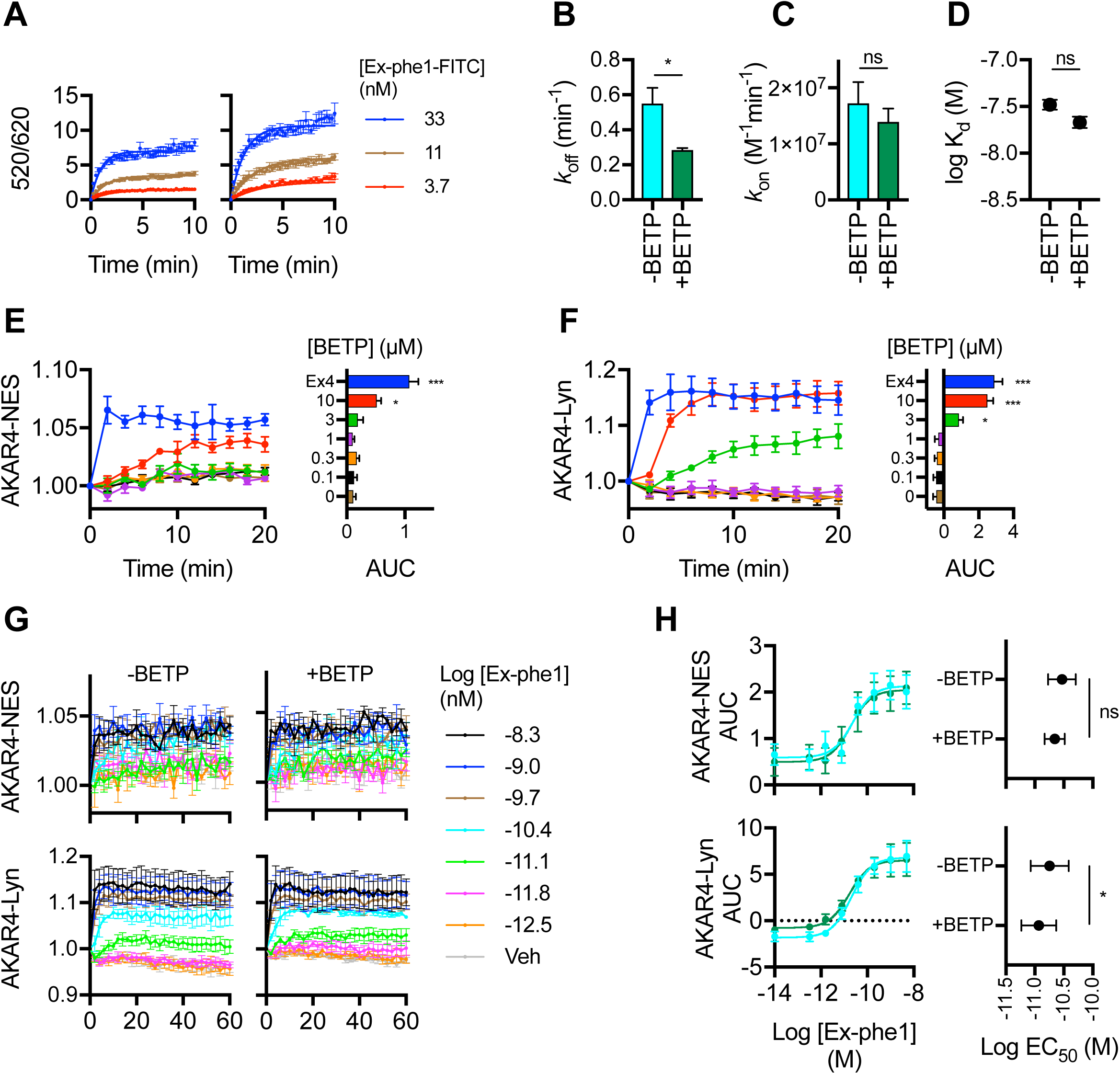
Effect of BETP on exendin-phe1 binding affinity and nanodomain signaling – extra data. (**A**) Kinetic binding response of exendin-phe1-FITC (Ex-phe1-FITC) with or without co-application of BETP (10 µM), measured by TR-FRET in Lumi4-Tb-labeled HEK293 cells stably expressing SNAP-GLP-1R, expressed ratiometrically as 520 / 620 nm signal, *n*=4. (**B**) Dissociation rate constant (*k*_off_) determined from (A), paired *t*-test. (**C**) Association rate constant (*k*_on_) determined from (A), paired *t*-test. (**D**) Equilibrium binding constant (K_d_) determined from (A), paired *t*-test. (**E**) Effect of BETP at the indicated concentration, or exendin-4 (Ex4, 100 nM) on AKAR4-NES FRET responses in monoclonal SNAP-GLP-1R cells, *n*=3, AUC calculated relative to individual normalized baseline, randomized block one-way ANOVA with Dunnett’s test *vs.* vehicle. (**F**) As for (E) but with AKAR4-Lyn. (**G**) Effect of co-application of BETP (1.5 µM) on AKAR4-NES and AKAR4-Lyn responses to exendin-phe1 in monoclonal CHO-SNAP-GLP-1R cells, expressed ratiometrically and after normalization to individual well baseline, *n*=3. (**H**) Dose-response curves constructed from AUC measurements of data shown in (G), 3-parameter fit shown, comparison of Log EC_50_ by paired *t*-test. Data are depicted as mean ± SEM; ns=non-significant, *p<0.05, ***p<0.001, by statistical test indicated in the text.

**Figure S7.**
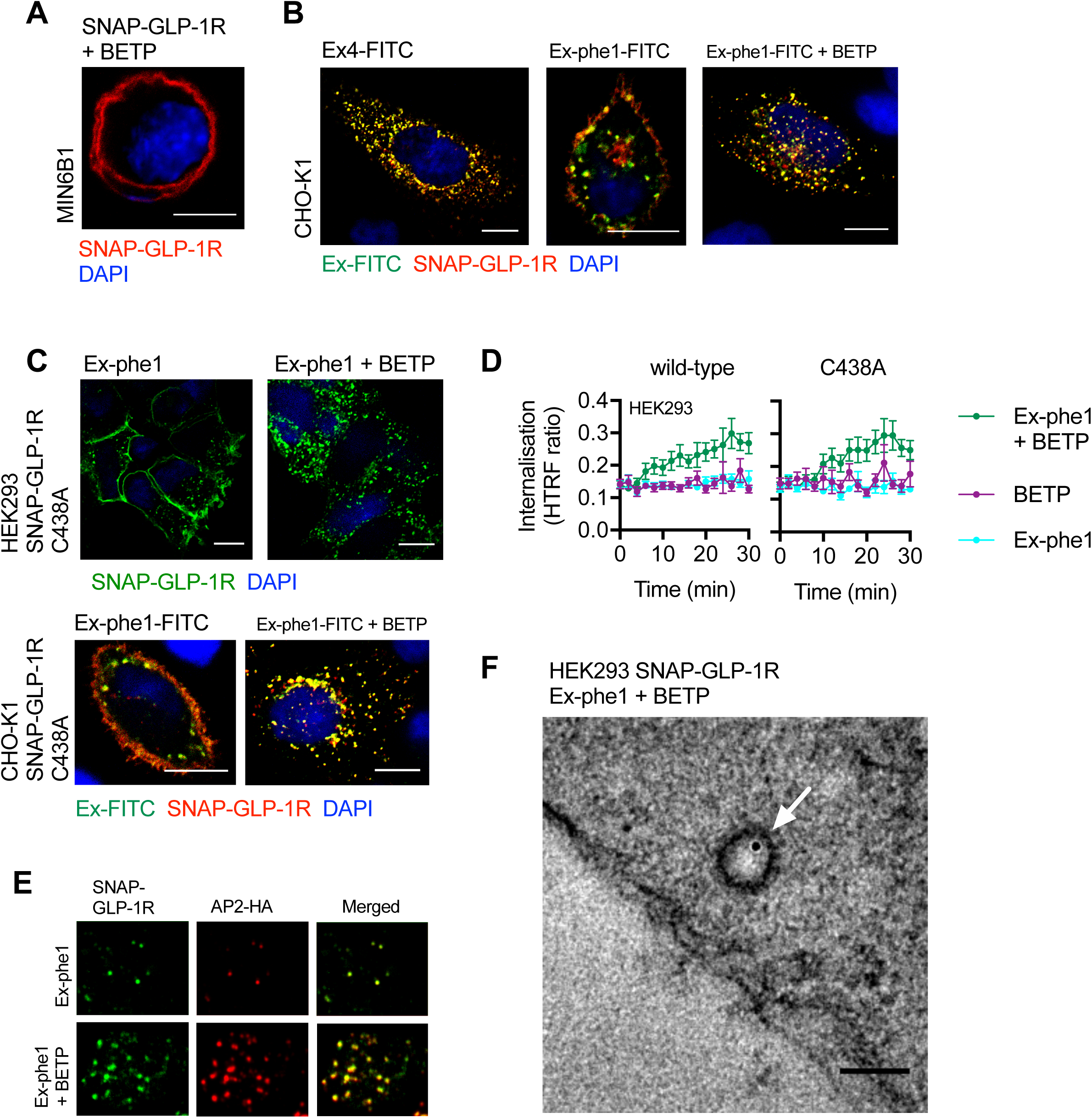
Effect of BETP on exendin-phe1 trafficking – extra data. (**A**) Effect of BETP (10 µM) on SNAP-GLP-1R localization in MIN6B1 cells stably expressing SNAP-GLP-1R. SNAP-Surface 549, red; nuclei (DAPI), blue; size bar, 10 µm. (**B**) Confocal analysis of FITC-agonist (green) and SNAP-GLP-1R (red) internalization in CHO-K1 cells stably expressing SNAP-GLP-1R following stimulation with 100 nM of the indicated FITC-agonist with or without BETP (10 µM) for 10 min. Nuclei (DAPI), blue; size bars, 10 µm. (**C**) Confocal analysis of SNAP-GLP-1R C438A localization in HEK293 (top) or CHO-K1 (bottom) cells following stimulation with exendin-phe1 (Ex-phe1, top) or exendin-phe1-FITC (Ex-phe1-FITC, bottom), with or without BETP (10 µM) for 10 min. SNAP-Surface 488, green (top); Ex-phe1-FITC, green and SNAP-Surface 549, red (bottom); nuclei (DAPI), blue; size bars, 10 µm. (**D**) SNAP-GLP-1R internalization measured by DERET in HEK293 cells transiently expressing wild-type or C438A mutant SNAP-GLP-1R, treated with exendin-phe1 (100 nM), BETP (10 µM), or both, expressed ratiometrically, *n*=6. (**E**) TIRF microscopy analysis of plasma membranes from MIN6B1 cells stably-expressing SNAP-GLP-1R and transiently transfected with µ2-HA-WT, which codes for the µ2 domain of the clathrin adaptor AP2 fused to an HA-tag (AP2-HA), labeled with SNAP-Surface 488 (green) and stimulated for 1 min with 100 nM exendin-phe1 with and without BETP (10 µM), prior to fixation and HA-tag immunofluorescence (red). (**F**) Electron micrograph depicting a representative image of a clathrin-coated pit-localized gold-labeled SNAP-GLP-1R (white arrow) in HEK293 cells stably expressing SNAP-GLP-1R stimulated for 1 min with 100 nM exendin-phe1 + BETP (10 µM); size bar, 100 nm. Data are depicted as mean ± SEM.

